# Phased epigenetic fragility and stability accelerate the genetic evolution of B-cells

**DOI:** 10.1101/2025.06.24.661408

**Authors:** Mark Y Xiang, Haripriya Vaidehi Narayanan, Vaibhava Kesarwani, Tiffany Wang, Alexander Hoffmann

## Abstract

Non-genetic heterogeneity can provide population robustness when responding to threats or making developmental decisions, but when the biological process rests on specific individuals (tissue-resident macrophages or genetic evolution), non-genetic heterogeneity degrades performance. Vaccine responses depend on the Darwinian evolution of B-cells to generate high-affinity, genetically-encoded antibodies, yet B-cell decision-making is non-genetically variable. In fact, B-cell epigenetic states are fragile during selection but stable during the proliferation burst. Here we report that the iterative dynamics of epigenetic fragility and stability of B-cell fate decisions do not impair but accelerate affinity maturation, by modulating the generation and removal of high-affinity outliers on a fitness landscape that is non-monotonic defined not only by survival/proliferation but plasma cell differentiation. These insights reconcile classical B-cell clonal selection theory with experimentally observed dynamics of epigenetic variability. The resulting model correctly predicts emergent vaccine response properties in alternate mouse strains and may contribute to personalized vaccination strategies.

**HIGHLIGHTS:**

- B-cell epigenetic states are fragile during antigen selection but stable within proliferative bursts
- Epigenetic state fragility allows high affinity cells to escape plasma cell differentiation
- Epigenetic state stability within proliferative bursts maximizes progeny of fittest cells
- Phased epigenetic dynamics of B-cells predicts emergent vaccine response properties

**GRAPHICAL ABSTRACT:** 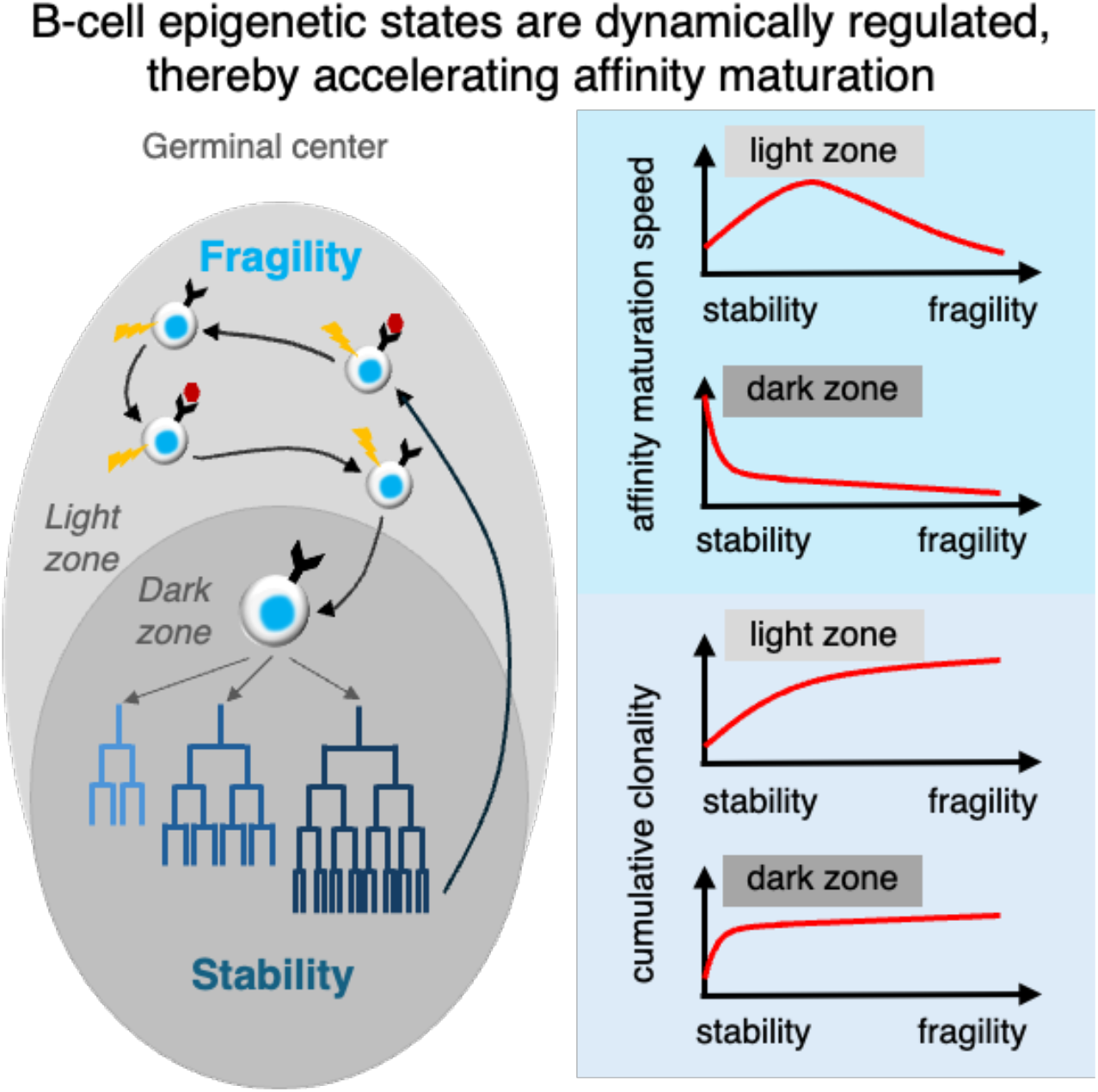

## INTRODUCTION

With the identification of DNA as the carrier of genetic information that provides for the heritable traits of organisms ^1^, Conrad Waddington coined the term epigenetics to refer to the set of instructions that act on top of genetic instructions in order to explain how distinct cell types with identical genetic information may arise during development ^2^. These instructions are heritable within a cell lineage but may gradually change or bifurcate during differentiation, thereby forming the Waddington landscape ^3^ Yet recent studies have revealed that the set of epigenetic instructions - contained in cells’ chromatin states ^4^, molecular regulatory networks ^5^, and subcellular organization ^6^ – exhibit substantial heterogeneity even among genetically identical cells of the same cell type. Unlike genetic mutations, which cause stable changes to the genetic instructions, epigenetic states are dynamic and may respond to environmental cues and molecular stochastic fluctuations. For example, following inheritance of the same epigenetic state, sibling cancer cells rapidly diverge in their propensity to respond to a death-inducing ligand ^7^. This epigenetic variability allows genetically identical cells to respond differentially to noxious environmental perturbations and may generate population robustness *via* a process called bet hedging ^8^.

However, in physiological processes that depend on reliable cell-decision-making by an individual, epigenetic variability may be detrimental. For example, the stimulus-response specificity of tissue resident sentinel cells is diminished by variable responses to defined stimuli ^9,10^. Further, in Darwinian processes that aim for the selection of genetic instructions that confer greatest fitness, epigenetic variability is likely detrimental ^11,12^. The generation of high affinity antibody by B-cells involves a Darwinian process of mutation and selection as described by classical clonal selection theory ^13^. Thus, the discovery of seemingly stochastic B-cell responses to identical stimulation was notable ^14,15^ and prompted the development of population dynamics models that assumed stochastic decision making ^16,17^. Pedigree lineage tracing through microscopy revealed that the epigenetic state was actually remarkably stable within the stimulus-induced proliferative burst and that the proliferative heterogeneity was largely due to epigenetic heterogeneity of the starting population ^18^. Yet, it remains unknown how this dynamically regulated epigenetic variability affects the Darwinian evolutionary process of selecting and producing high affinity antibodies.

In response to vaccination, B cells migrate between two compartments within the lymph node’s germinal center (GC): the light zone (LZ), where they undergo selection by picking up antigen and identifying cognate T-cells to provide pro-proliferative signals, and the dark zone (DZ), where they proliferate over several divisions and mutate their antibody genes before cycling back to the LZ. Over a two-week vaccine response time there are estimated to be up to 40 GC cycles of selection and proliferation/mutation ^19,20^. Moving around the LZ, B-cells, while in the G1 cell cycle phase, encounter numerous signals that impinge on their molecular networks, that may generate a highly heterogeneous set of epigenetic states. In contrast, B-cells in the DZ proliferate rapidly *via* a shortened S/M phase cell cycle and largely preserve their epigenetic state. Thus, the GC reaction is characterized by two phases that repeat for each GC cycle: a phase of “epigenetic fragility” and a phase of relative “epigenetic stability”.

Here, we addressed the question of how the dynamic control of epigenetic variability of B-cells impacts the antibody vaccine response. As there are no experimental tools to manipulate epigenetic variability in defined ways, we developed a mathematical model of the antibody maturation process in the GC by leveraging a wealth of experimental data before validating it with perturbation studies in mouse mutants. This mathematical model encodes key aspects of classical clonal selection theory, experimental measurements of cell numbers, GC characteristics, mutation rates, selection stringency, as well as the heterogeneous cell fate decision processes of cell death/survival, proliferation, and differentiation. By manipulating the generation of epigenetic fragility (“stochasticity”) or its stability (“heritability”) we discovered how the phased fragility and stability of B-cell epigenetic states does not impair but accelerates the evolution of high affinity B-cells.

## RESULTS

### A knowledge-based mathematical model of B-cell affinity maturation

To study how non-genetic heterogeneity of B-cells affects their genetic evolution that drives antibody maturation, we leveraged extensive prior knowledge ^21–29^ to construct a mathematical model of the Darwinian process of B-cell selection and clonal expansion in germinal centers (GCs) (Fig. 1A). We parametrized the initial B-cell population size based on the number of clones seeding a typical GC ^30^, starting with a distribution of low antigen affinities ^31^. We abstracted the spatial segregation of the GC into light and dark zones and B-cell migration between them into sequentially occurring processes within each zone. The model incorporates repeated cycles of three core processes: affinity-dependent selection in the light zone, followed by stimulus-dependent cell fate decisions to survive, proliferate, or differentiate in the dark zone, with mutation during proliferative clonal bursts.

**Figure 1.**
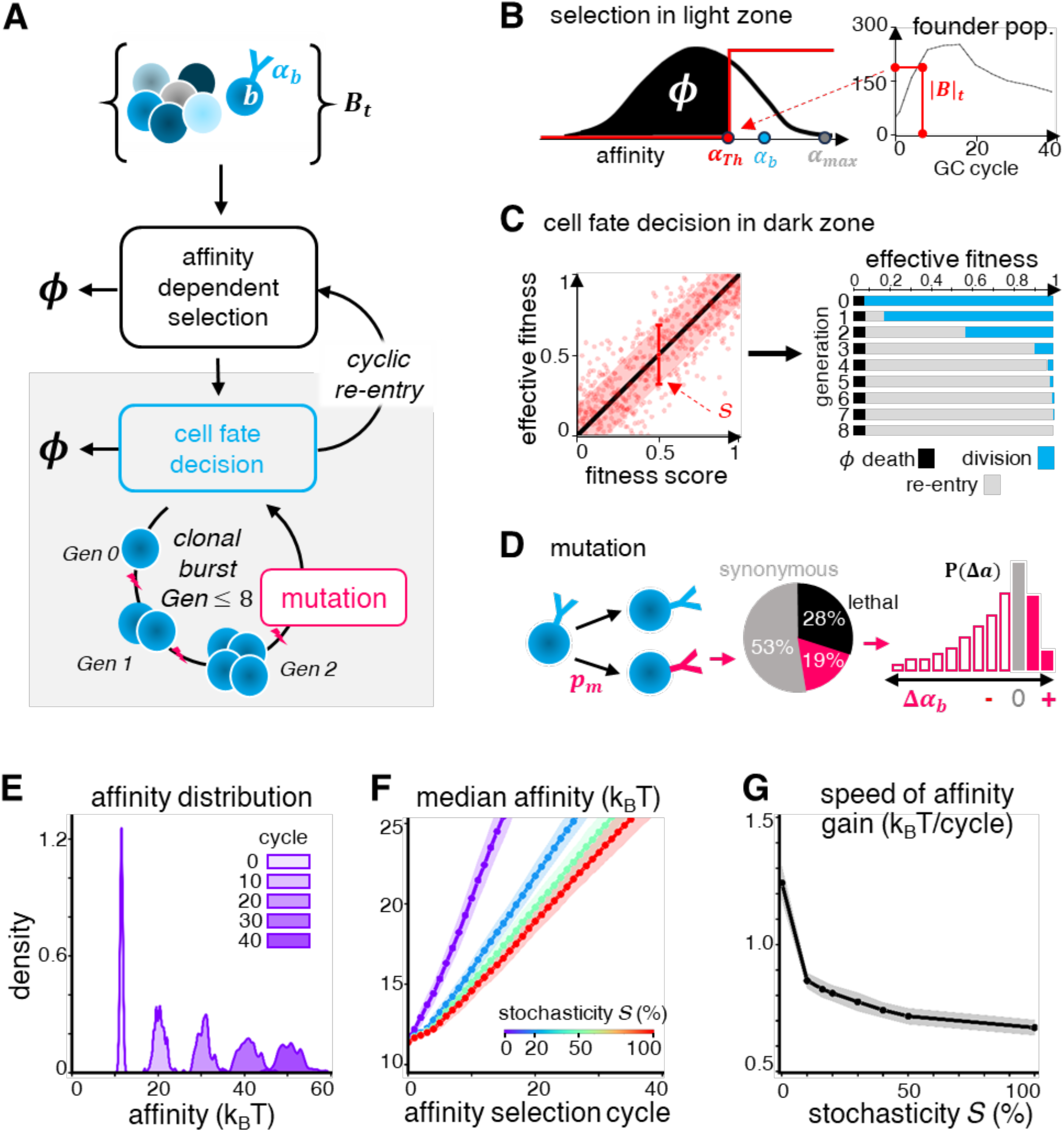
A mathematical model of the Darwinian process underlying antibody generation suggests that stochastic B-cell fate decisions are detrimental to affinity gain. **(A)** Schematic of the mathematical model of affinity maturation incorporating stochasticity in cell fate decision making ^21–31^. The model includes sequential processes of affinity-dependent selection in the light zone and cell fate decisions in the dark zone (boxed grey) across multiple affinity maturation cycles, with mutation occurring during proliferative bursts in the dark zones. *B*_*t*_ represents the population of B-cells at any given cycle, and ϕ represents the removal of B-cells due to death. **(B)** Schematic of parameterizing affinity-dependent selection in the light zone. Based on the size of the germinal center (right), we define an affinity threshold *α*_*th*_ which selects a given number of highest affinity B-cells. This threshold varies dynamically across cycles, due to changes in both GC size and the B-cell affinity distribution ^34^. **(C)** Schematic of parameterizing affinity-dependent but stochastic cell fate decisions in the dark zone. **(C – Left)** Stochastic cell fate decisions are modeled using a stochasticity parameter *S* to distribute affinity-dependent fitness scores. **(C – Right)** The resulting effective fitness scores are used to determine the corresponding B-cell fate decisions using a generation-specific probabilistic cell fate map. Higher fitness scores correspond to more proliferation while lower fitness scores favor cell death, consistent with affinity-dependent clonal selection theory ^13,44^. **(D)** Schematic of parameterizing mutation and affinity changes during proliferative bursts in the dark zone. During each cell division, each daughter cell independently mutates its B-cell receptor with a probability *p*_*m*_, set at 0.5 based on literature ^39,40^. The pie chart indicates the proportion of synonymous (grey), lethal (black), and non-synonymous (magenta) mutations ^41^. For the fraction of non-synonymous mutations, the histogram represents the probability of changes in B-cell affinity, where grey indicates no change, open bars detrimental mutations, and shaded bars affinity-enhancing mutations ^42^. **(E)** Kernel density plots showing the distribution of affinities of all B-cells re-entering the subsequent GC selection cycle, at regular intervals during affinity maturation when cell fate stochasticity is set to zero. Note the heavy tail which contains cells driving the affinity gain through the Darwinian process. **(F, G)** Line plots of **(F)** the median affinity and **(G)** the speed of affinity gain (defined as the increase in median affinity relative to the previous selection cycle) for B-cells re-entering each subsequent affinity maturation cycle, across varying degrees of cell fate stochasticity. Points indicate the median and shaded regions indicate the 95% confidence interval across 100 simulation runs.

During affinity-dependent selection, either the initial founder B-cells, or their progeny re-entering subsequent cycles after a clonal burst, are positively selected based on a dynamic affinity threshold ^32^ (Fig. 1B left). This is a result of competition for limited T-cell help ^33^, and shifts with time ^32^ reflecting the carrying capacity of the GC, which expands in size before gradually contracting ^34^ (Fig. 1B right). Low-affinity B-cells undergo apoptosis, while high-affinity B-cells proceed to cell fate decisions.

In contrast to models that assign stochastic B-cell fates based on expected outcome ^24,25,35^, this model assigns affinity-dependent fate decisions as prescribed by classical clonal selection theory ^13^. For each B-cell that is positively selected, a fitness score is assigned corresponding to the rank of its affinity in the population, as a proxy for signaling by T-follicular helper cells. Fitness scores are then mapped onto cell fate decisions described by a multi-generational cell fate map that conforms to classical clonal selection theory ^36^ and literature measurements ^37,38^ (Fig. 1C right, details depicted in Fig. S1A). To capture the non-genetic heterogeneity in B-cell proliferation, we introduced a stochasticity parameter *S* that describes the degree of decorrelation between the initial affinity-dependent fitness score and the actual cell fate decisions made in successive generations of a proliferative burst (Fig. 1C left, details in Fig. S1A).

To complete the Darwinian process we modeled the generation of genetic variants: B-cells undergoing division activate somatic hypermutation, which alters the receptor’s affinity for the antigen based on the probability of mutation occurrence ^39–41^ and the probability that the mutation results in a change in affinity versus being detrimental (cell death) or neutral (no change in affinity or fitness) ^42^ (Fig. 1D). We ensured reproducibility in the probabilistic formulations of cellular behaviors by performing sufficient Monte Carlo simulations to achieve 95% confidence within a 1% margin of error of the median estimate. First simulations of the model showed, as expected for a Darwinian process, that antibody affinity increases over time (Fig. 1E), demonstrating the capacity to simulate affinity maturation as a function of the key regulatory steps described in clonal selection theory when parameterized by a wealth of published experimental data. Affinity distributions were characterized by a tail of high-affinity outliers that becomes more pronounced at later cycles (Fig. 1E), indicating that affinity maturation is an outlier-driven process.

### Simplest model: Stochasticity in B-cell-fate decisions impairs affinity gain

Prior *in vitro* studies of B-cell proliferation revealed a high degree of non-genetic cell-to-cell heterogeneity in proliferative responses ^15,16^ such that probabilistic cell fate decision models could capture the heterogeneous population dynamics ^43^. With the newly established model we could now investigate the impact of this probabilistic cell fate decision process on antibody affinity maturation. Simulations showed that with increasing stochasticity *S*, the gain in median affinity is dramatically reduced (Fig. 1F), as the tail of high-affinity outliers in later cycles is diminished (Fig. S1B). Quantifying the rate of affinity gain as a function of the stochasticity parameter confirmed this conclusion (Fig. 1G).

Thus, the simple model of Darwinian evolution predicted that the generation of epigenetic heterogeneity in the propensity of cell fate decision making, a fundamental hallmark of the biology of cells, would substantially impair the speed of affinity maturation. However, B-cell biology in the GC reaction involves two additional fate decision events: differentiation into memory B-cells and antibody-producing plasma cells.

### Considering memory B-cell differentiation: Stochasticity improves clonal diversity

Low-affinity B-cells give rise to memory B-cells which may seed future antigen responses, while high-affinity B-cells differentiate into antibody-secreting plasma cells ^44,45^. We refined the model by incorporating differentiation into memory B-cells or plasma cells as additional cell fate decisions (Fig. 2A-C). This allows us to contrast the removal of GC B-cells from the Darwinian evolutionary process at either end of their affinity distribution.

**Figure 2.**
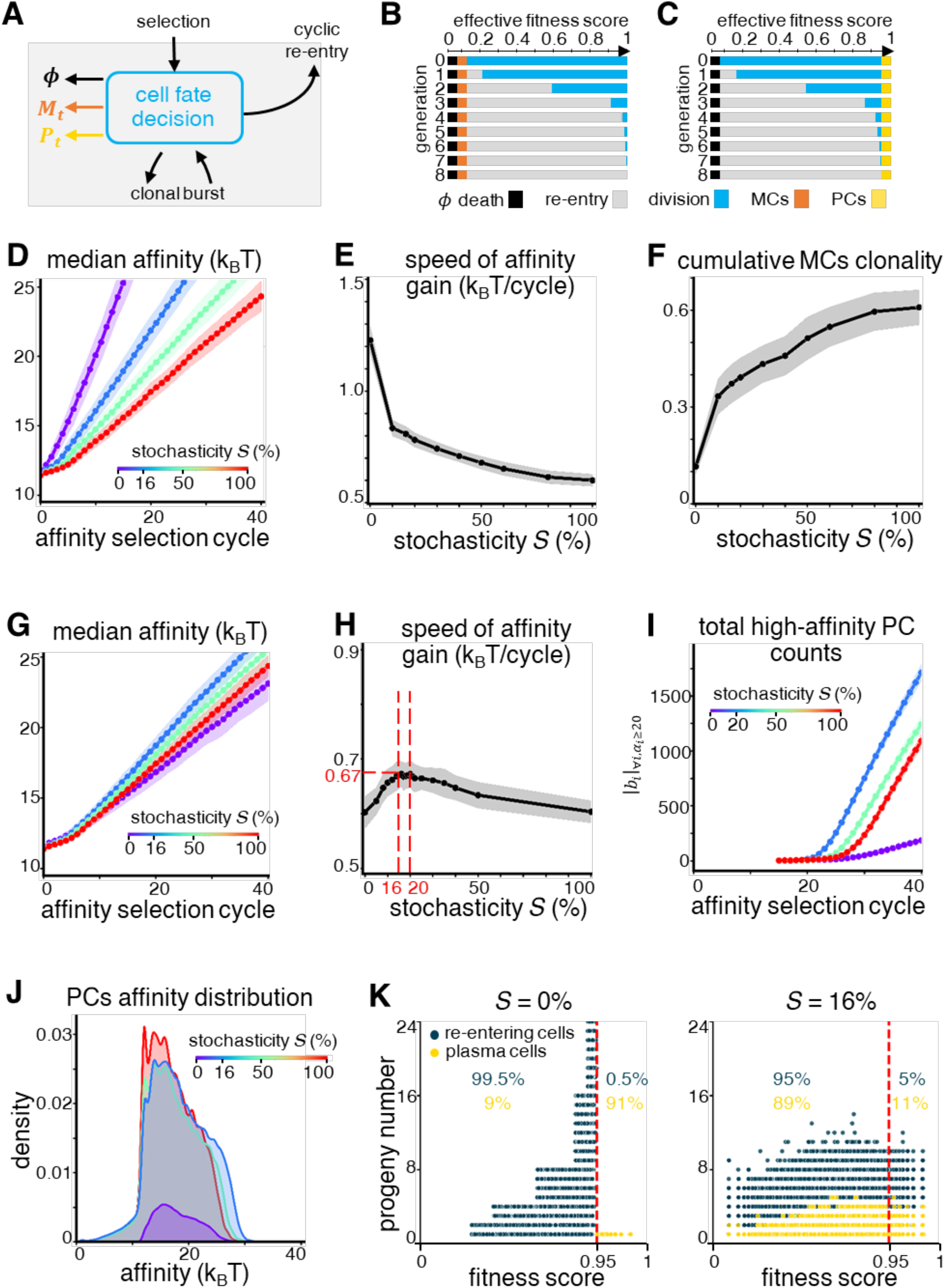
Considering memory and plasma cell differentiation, an optimal degree of stochasticity in B-cell fate decisions emerges. **(A)** Schematic of modified cell fate decision module incorporating memory B-cell *M*_*t*_ and plasma cell differentiation *P*_*t*_ ^13,44^. **(B)** Schematic of the modified cell fate decision map, directing the 5% of B-cells just above the lowest effective fitness scores (undergoing cell death) in each generation to differentiate into memory B-cells ^23,49^. **(C)** Schematic of the modified cell fate decision map, directing B-cells with the top 5% of effective fitness scores in each generation to differentiate into plasma cells ^23,49^. As in Figure 1C, effective fitness scores of founders and progeny in each cycle are distributed as specified by the stochasticity parameter S. **(D-F)** Line plots of **(D)** the median affinity of cycling GC B-cells re-entering the light zone for a subsequent round of selection, **(E)** their speed of affinity gain (defined as the increase in median affinity relative to the previous selection cycle), and **(F)** the cumulative clonal diversity of memory B-cells (defined as fraction of extant clones relative to number of founder B-cell clones seeding the germinal center at cycle 0) generated in each affinity maturation cycle, for varying degrees of cell fate stochasticity. Points indicate the median and shaded regions indicate the 95% confidence interval across 100 simulation runs. **(G-I)** Line plots of **(G)** the median affinity of cycling GC B-cells re-entering the light zone for a subsequent round of selection, **(H)** their speed of affinity gain (defined as the increase in median affinity relative to the previous selection cycle), and **(I)** the total number of plasma cells with high affinity (defined as above 20.7 *k*_*B*_*T* or *K*_*D*_ 1 nM) generated in each affinity maturation cycle, for varying degrees of cell fate stochasticity. Points indicate the median and shaded regions indicate the 95% confidence interval across 100 simulation runs. The vertical red lines in **(H)** indicate the region where speed of affinity gain is highest, between 16% to 20% stochasticity. **(J)** Kernel density plots showing the distribution of plasma cell affinities generated at different degrees of cell fate stochasticity. **(K)** Progeny plot showing the number of GC-reentering cells (blue dots) and plasma cells (yellow dots) that derive from a single progenitor whose affinity-dependent fitness score is shown on x-axis (cycle 21). The red line indicates the 5% fitness score cut-off for plasma cell differentiation ^23,49^. Percentages indicate the fraction of re-entering cells (blue) and plasma cells (yellow) that are below and above this fitness score cut-off.

While the lowest affinity B-cells are eliminated by death, we modeled the next 5% of B-cells at low affinity differentiating into memory B-cells ^46–48^ (Fig. 2B). We explored how stochasticity in cell fate decisions affects affinity maturation when low-affinity variants are removed. As earlier, we found that increasing stochasticity substantially reduces affinity gain of cycling GC B-cells (Fig. 2D, E), and hence memory B-cells sampled from this pool (Fig. S2A). However, stochasticity dramatically improves the clonal diversity of memory B-cells (Fig. 2F) and their total numbers produced (Fig. S2B), though at the cost of mutational depth (Fig. S2C).

### Considering plasma cell differentiation: Stochasticity enhances affinity gain

We further revised the cell fate map based on literature measurements ^23,37,38,49^ with 5% of B-cells at the highest affinities expected to differentiate into plasma cells (Fig. 2C). We again explored how stochasticity in cell fate decisions affects gains in the affinity of plasma cells.

Increasing the stochasticity parameter from 0% to 100% revealed that the relationship between stochasticity and the affinity maturation rate is no longer monotonic when high-affinity outliers are removed as plasma cells (Fig. 2G). Specifically, we observed an optimal level of stochasticity (between 15-25%, peaking at ∼16%) at which cycling GC B-cells showed the fastest affinity maturation speed (Fig. 2H). We confirmed this by analyzing the affinity distribution of plasma cells, which displayed similar trends in maturation dynamics (Fig. S2D, E).

Our results indicated that similarly to memory B-cells, stochasticity in cell fate decisions dramatically increases the total number of plasma cells (sampled from the large pool of GC B-cells), and this effect persists even at high levels of stochasticity (Fig. S2F). Likewise, increasing stochasticity also improved clonal diversity of plasma cells (Fig. S2G, H), while diminishing their mutational depth (Fig. S2I). However, plotting plasma cell affinities revealed that only the moderate level of stochasticity (16%) yields the largest proportion of high-affinity plasma cells (Fig. 2I), i.e. increases the tail of high-affinity outliers in the distribution of their affinities (Fig. 2J). This suggests that moderate stochasticity may optimize affinity maturation by balancing plasma cell differentiation and progeny production among high-affinity cells.

To further investigate how stochasticity could result in a gain in affinity maturation, we examined the progeny of selected cells that are either differentiating into plasma cells or re-entering the germinal center for another evolutionary cycle (Fig. 2K). At 0% stochasticity, all highest affinity cells differentiate into plasma cells without any further proliferative expansion, while almost all cells re-entering selection (99.5%) are below the high-affinity threshold. However, at the optimal 16% stochasticity, there is a 5% tail of high-affinity cells that complete the proliferative program to enter another round of selection and may therefore contribute to subsequent cycles of selection and mutation. Also, while only 11% of plasma cells now cross the high-affinity threshold, their total numbers have been amplified by division prior to differentiation.

In other words, stochasticity in cell fate decisions allows some high affinity cells to escape removal from the Darwinian process, thereby accelerating affinity maturation gains. Yet, stochasticity also allows lower affinity B-cells to become plasma cells, reducing the stringency of affinity-dependent differentiation. Thus, as stochasticity increases, its detrimental impact on affinity-based selection dominates over the benefit. This results in an optimal level of stochasticity in cell fate decisions for efficient affinity maturation that improves both affinity and clonal diversity of plasma cells.

### Within-burst heritability of epigenetic states stabilizes proliferative fitness

Following selection in the light zone, B-cells undergo a clonal burst of rapid, successive divisions over multiple generations in the dark zone. Live cell microscopy of stimulus-induced B-cell proliferation revealed that sibling and cousin cells make highly correlated cell fate decisions, suggesting that the wide heterogeneity of B-cell responses to selection signals (*via* BCR and CD40) are in fact based on heterogenous epigenetic states in selected cells that are largely inherited by the progeny within a clonal burst ^18^. The stability of these epigenetic states of proliferating B-cells is in contrast to their relative instability in a cancer cell line, where correlations between cell death decisions of siblings are lost within hours ^7^.

Expanding the model to account for within-burst heritability of the non-genetic cell state, we examined how it might shape B-cell proliferation dynamics in the GC and the maturation of antibody affinity. Specifically, we introduced a heritability parameter that reduces stochasticity across successive generations within a clonal burst, allowing us to study its effects (Fig. 3A). We could thereby generate lineage trees with and without within-burst heritability. Without heritability of the non-genetic cell state, progeny cells made independent fate decisions, generating asymmetric trees (Fig. 3B top). In contrast, high heritability (95%) led to a more cohesive symmetric lineage structure (Fig. 3B bottom), where sibling cells made highly correlated cell fate decisions in line with experimental observations ^18^.

**Figure 3.**
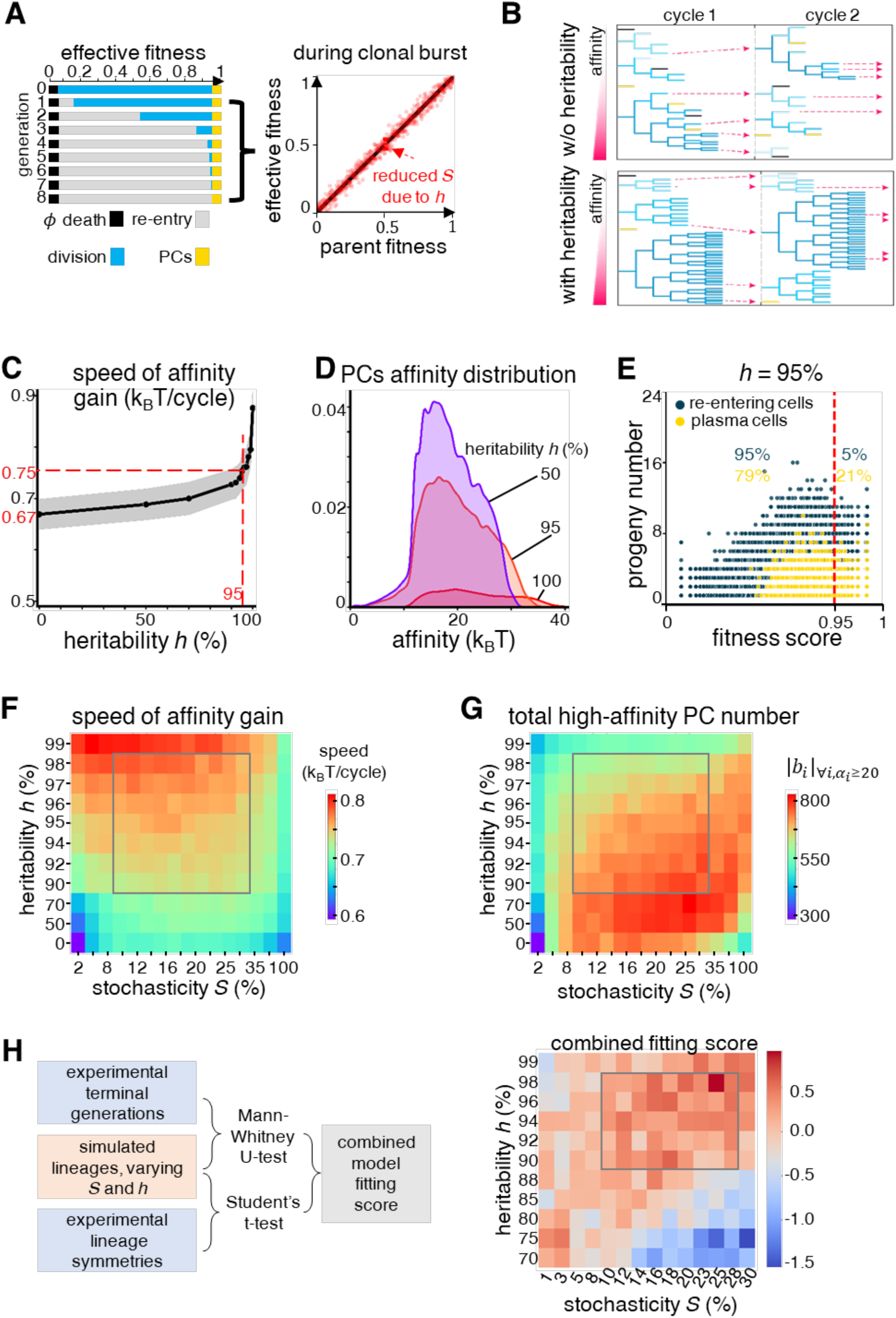
Phased stochasticity and heritability of B-cell fate decisions enables both rapid affinity gains and high plasma cell counts. **(A)** Schematic showing generation-specific cell fate map (left) with reduced stochasticity *S* due to heritability *h* (right, due to heritability) for progeny in Generation 1 onwards of a proliferative burst as observed by live cell microscopy ^18^. **(B)** Example lineage trees across two successive selection cycles, at optimal 16% stochasticity modified by a heritability parameter of 0% (top) or 95% (bottom) within proliferative bursts. In each generation, dead cells are indicated in black, proliferating cells in shades of blue, and plasma cells in yellow. Pink arrows show cells re-entering the next selection cycle. **(C)** Line plot of the speed of affinity gain (defined as the difference between median plasma cell affinities generated in the current and previous selection cycles), for optimal 16% cell fate stochasticity among founder B-cells but varying degrees of heritability during proliferative bursts. Points indicate the median and shaded regions indicate the 95% confidence interval across 100 simulation runs. The red lines indicate the rise in speed of affinity gain from 0% to 95% heritability. **(D)** Kernel density plots showing distribution of plasma cell affinities generated at different degrees of heritability and optimal 16% cell fate stochasticity. **(E)** Progeny plot showing the number of GC-reentering cells (blue dots) and plasma cells (yellow dots) that derive from a single progenitor whose affinity-dependent fitness score is shown on x-axis (cycle 21). The red line indicates the 5% fitness score cut-off for plasma cell differentiation ^23,49^. Numbers indicate the fraction of re-entering cells (blue) and plasma cells (yellow) that are below and above this fitness score cut-off. **(F-G)** Two-dimensional heat map of **(F)** the speed of affinity gain and **(G)** total number of plasma cells above the high-affinity threshold of 20.7 *k*_*B*_*T* (corresponding to *K*_*D*_ 1 nM) ^31^ across varying degrees of both stochasticity and heritability in B-cell fate decisions. **(H – Left)** Workflow for estimating the stochasticity and heritability of B-cells using experimental data of tracked single WT B-cell lineages from proliferative bursts *in vitro* ^18^. **(H – right)** Heatmap of the resulting fitting scores across stochasticity and heritability, with higher scores indicating better fits to the experimental data. In **(F-H)**, every heat map pixel represents the median of 100 simulation runs (300 runs for **(H)**), and the gray boxes outline the physiologically plausible range of values for stochasticity and heritability based on experimental observations ^18^.

We then explored how within-burst heritability in the model impacts affinity maturation in the plasma cell population, at the previously identified optimal value of 16% stochasticity. Unlike the stochasticity in cell fate decisions, within-burst heritability of fate decision propensities showed a monotonic but non-linear gain in affinity maturation, accelerating affinity gain (Fig. S3A). The speed of affinity gains increased dramatically within the biologically relevant range (Fig. 3C), suggesting an evolutionary benefit for maintaining the non-genetic cell state during a clonal burst.

Our results also indicated that high within-burst heritability reduced the total number of plasma cells (Fig. S3B), while promoting higher affinity (Fig. S3C), as observed in the plasma cell affinity distribution (Fig. 3D). This implies a trade-off between plasma cell production and affinity gain. To further understand this, we accessed progeny counts of founder cells, distinguishing between those differentiating into plasma cells and those re-entering the GC light zone. We found that high heritability restricts the affinity threshold for plasma cell formation, increasing the proportion of plasma cells above the high-affinity threshold, from 11% high-affinity plasma cells without any heritability to 21% high-affinity plasma cells at 95% heritability (Fig. 3E).

### Dynamic regulation epigenetic fragility and heritability optimizes affinity maturation

We extended our analysis to examine the combined effects of stochasticity in the cell fate decision and within-burst heritability. We found that the greatest speed of affinity gain occurs at around 16% stochasticity and high heritability levels (95% to 99%, Fig. 3F), consistent with results from individual parameter analyses (Figs. 2D, 3C). As expected, at a fixed stochasticity or heritability value, varying the other parameter confirmed previous findings. However, the number of high-affinity plasma cells peaked at a different optimum, with similar stochasticity (around 16% to 25%) but intermediate values of heritability (50% to 70%, Fig. 3G), reiterating that heritability modulates a trade-off between plasma cell numbers and affinities.

We then evaluated how the predicted optima for stochasticity and heritability correspond with experimental observations (Fig. 3H). We estimated stochasticity in proliferative bursts by the spread of terminal generations observed in dye dilution assays ^15,18^, and heritability by lineage symmetry in single-cell live microscopy assays ^18^ under identical stimulus conditions. By iteratively comparing measured distributions of terminal generations to model simulations that independently varied both parameters, we estimated the best-fit at moderate stochasticity between 12% to 28% and high heritability between 90% to 98%, consistent with the predictions for maximizing affinity gain.

We also compared the effects of stochasticity (Fig. S2C-F) and heritability (Fig. S3D-G) on clonality and mutational depth of plasma and memory cells, as additional dimensions of the GC response. We found that heritability had a modest but opposite effect to stochasticity. At higher heritability, increased progeny within a given lineage led to more mutations, while restricting the number of clones selected at a given carrying capacity.

Taken together, epigenetic heterogeneity among founder B-cells and its within-burst heritability balance multi-dimensional properties of speed of affinity gain, titer, clonality, and mutational depth, defining a biological set-point for the antibody response. The congruence between predicted and experimentally fitted stochasticity and heritability parameters underscores the role of phasing epigenetic fragility and stability in optimizing production and maturation of high-affinity plasma cells, supporting an effective immune response.

### Predicting and testing vaccine responses in distinct mouse strains

B-cell fate decision-making propensities are determined by molecular regulatory mechanisms which may differ between individuals due to genetics or health status, while epigenetic variability distributes these fate decisions. We asked whether our model incorporating the phasing of epigenetic variability could predict vaccination response outcomes based on B-cell fate decision propensities.

We experimentally tested model predictions of affinity maturation through perturbation studies using genetically modified mice. Specifically, we chose *nfkb1*^-/-^ and *IκBα*^*SS/SS*^ mouse strains as they alter the regulatory dynamics of the NFκB signaling pathway that drives B-cell fate determination. We undertook *in vitro* dye dilution studies to measure cell fate decision proportions (death, division, PC differentiation) across each generation of a proliferative burst (Fig. 4A) and used this to construct probabilistic cell fate maps for WT, *nfkb1*^-/-^ and *IκBα*^*SS/SS*^ mice (Fig. S4A-D). Our measurements showed that compared to WT, *nfkb1*^*-/-*^ mice have increased cell death but reduced plasma cell differentiation, whereas *IκBα*^*SS/SS*^ mice show increases in both.

**Figure 4.**
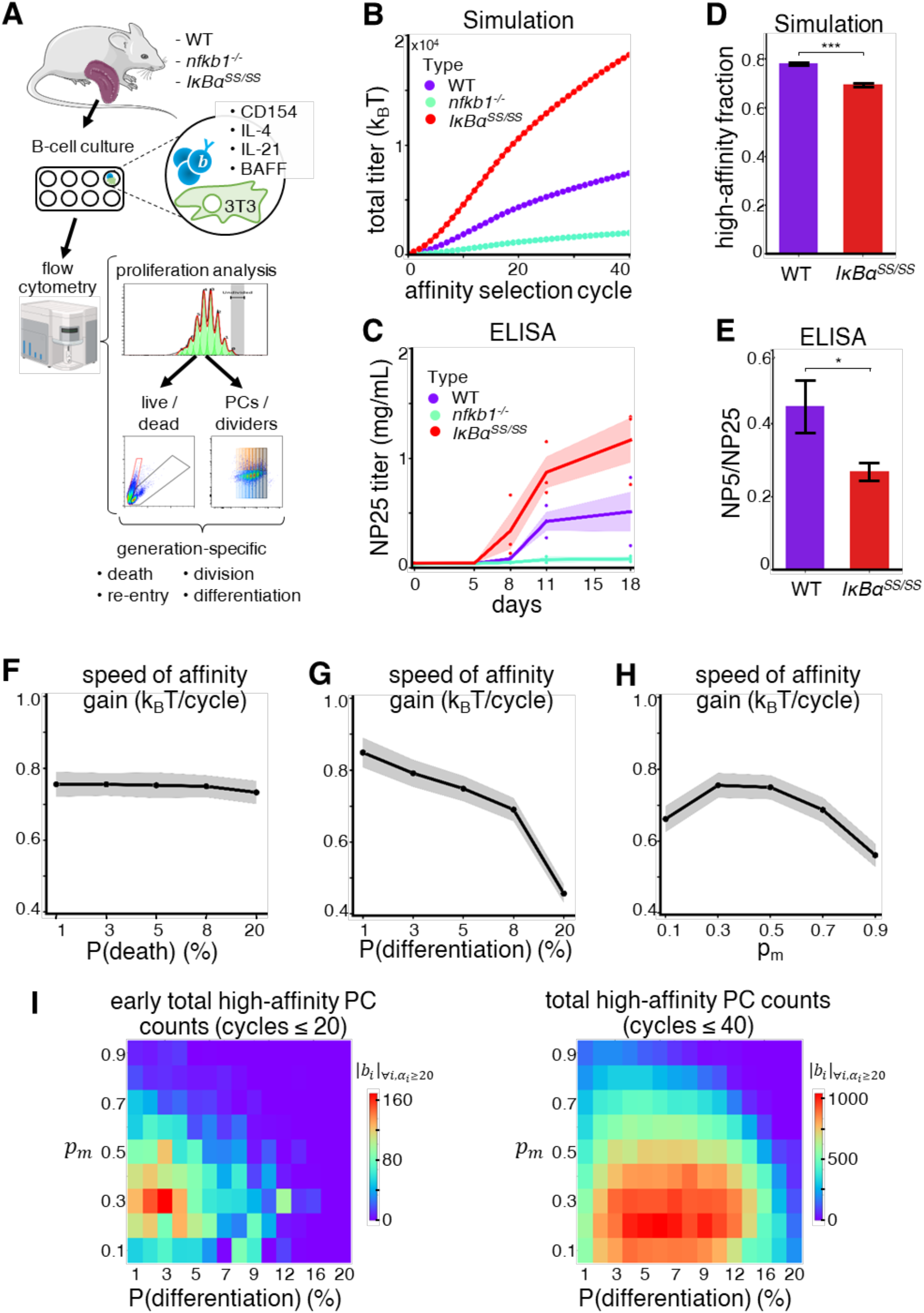
Key B-cell biological parameters in cell-fate decision-making determine trade-offs in vaccine-responsive affinity maturation outcomes. **(A)** Schematic of workflow to generate probabilistic cell fate maps from *in vitro* B-cell culture for three genotypes (WT, *nfkb1*^-/-^, *IκBα*^SS/SS^) varying in cell fate propensities. Follicular B-cells from naïve mouse spleens are cultured on a stromal layer of 3T3 fibroblasts with survival cytokines (BAFF) and a median (non-saturating) dose of T-dependent stimuli (CD154, IL-4, IL-21), to mimic the germinal center microenvironment while ensuring uniform stimulation and consistency with prior studies of proliferative heterogeneity. Proportions undergoing different cell fates within each generation are assessed at 120 hours by flow cytometry. **(B)** Model-predicted binding antibody titers (estimated as the cumulative plasma cell affinities) in each of the three genotypes, based on probabilistic cell fate maps generated from the *in vitro* flow cytometry measurements. Each point represents the median titer value across 100 simulation runs. **(C)** Line plots of *in vivo* serum antibody titers measured by ELISA against NP_25_ in mice from each genotype immunized with NP_21_-OVA. Each dot represents a measurement from one mouse, with solid lines showing the mean and shaded regions the standard error across three biological replicates for each genotype. **(D)** Bar chart of predicted high-affinity antibody fractions relative to total titer, estimated as the ratio of cumulative plasma cell affinities for those above 20.7 *k*_*B*_*T* (corresponding to *K*_*D*_ 1 nM) ^31^ versus the whole population, in WT and *IκBα*^SS/SS^ genotypes using measurements from day 11 to 18. Error bars represent standard errors across 100 simulation runs for each genotype. Differences between bars are statistically significant with p-value 10^−54^ by a Students’ T-test. **(E)** Measured high-affinity antibody fractions in WT and *IκBα*^SS/SS^ genotypes, evaluated as the ratio of antibody binding to NP_5_ (stringent, high-affinity titer) versus NP_25_ (total titer). Error bars show standard errors across 3 biological replicates per genotype. Differences between bars are statistically significant with p-value 0.037 by a Students’ T-test. **(F-H)** Line graphs showing how the speed of affinity gain (defined as the difference between median plasma cell affinities generated in the current and previous selection cycles) varies with key cell biological parameters representing the rates of **(F)** cell death, **(G)** plasma cell differentiation, and **(H)** receptor mutation. Points indicate the median and shaded regions indicate the 95% confidence interval across 100 simulation runs. **(I)** Heat maps of the total number of plasma cells generated above the high-affinity threshold (20.7 *k*_*B*_*T* corresponding to *K*_*D*_ 1 nM) ^31^, during the early (left) or the later phase (right) of the GC response. All outcomes in **(F-I)** are evaluated for the optimal 16% stochasticity and 95% heritability in cell-fate decision-making, and represent the median of 100 simulation runs.

Model simulations using these measured cell fate maps, together with previously fitted stochasticity and heritability parameter values, predicted higher total antibody titer for *IκBα*^*SS/SS*^ mice and lower for *nfkb1*^-/-^ mice compared to WT (Fig. 4B), consistent with their respectively higher or lower plasma cell differentiation rates (Fig. S4D). We confirmed these predictions by *in vivo* serum antibody titers in mice immunized with the model antigen NP_21_-OVA, measured by ELISA against NP_25_ to capture total NP-specific antibody (Fig. 4C). However, our simulations also predicted an unexpected decline in affinity maturation efficiency in *IκBα*^*SS/SS*^ mice (Fig. 4D). We confirmed this experimentally through diminished ratio of high-affinity (NP_5_-binding) to total (NP_25_-binding) antibodies in ELISA-based affinity ratio measurements (Fig. 4E). On the other hand, simulations using the measured cell fate maps but lacking either stochasticity (S = 0, corresponding to deterministic clonal selection theory) or heritability (S = 16%, h = 0, corresponding to the Cyton model) were unable to reproduce the *in vivo* observations (Fig. S4E, F).

These results predict emergent properties of affinity maturation *in vivo*, parametrized only on the underlying distribution of B-cell fate decisions measured *in vitro*. Further, incorporating the dynamic regulation of epigenetic variability is necessary to establish this robust alignment between knowledge-based mechanistic model predictions and experimental outcomes.

### Affinity maturation as a function of B-cell regulatory mechanisms

When underlying mechanisms are well-captured, the strength of mechanistic models lies in both predictive power as well as interpretability. To explain the observed relationships between B-cell fate decisions and vaccine response outcomes, we used the model to explore how the relevant regulatory mechanisms impact affinity maturation. We looked at the cell death rate which removes low-affinity variants and plasma cell differentiation rate which removes high-affinity variants to produce plasma cells, which varied between our mutant strains. We further studied the effect of mutation rate which controls the generation of these variants.

We found that death rate alone has little impact on affinity gain (Fig. 4F), plasma cell numbers and affinities (Fig. S5A, B). We attribute this to the exponential growth of high affinity B-cells over eight generations in a single GC cycle, rendering the system insensitive to the fate of lower-affinity cells. Extending this argument to memory B-cells, which differentiate from lower affinity cells, affinity gain and plasma cell numbers are unaffected. However, increasing this rate has clear benefits in improving clonal diversity of memory B-cells (Fig. S5C).

At the other end of the affinity range, increased plasma cell differentiation rate reduced speed of affinity gain (Fig. 4G), as this removes high-affinity B-cells from the Darwinian process. However, this also increases the number of effector plasma cells (Fig. S5D). The balance results in a trade-off between titer and affinity as a function of differentiation rate, with optimal production of high-affinity plasma cells at moderate rates between 3% to 8% (Fig. S5E). This explains why increased plasma cell differentiation produces higher titer but lower affinity of antibodies in *IκBα*^*SS/SS*^ mice compared to WT. Low-affinity B-cells and so memory B-cell clonality remain unaffected (Fig. S5F).

We further found that the mutation rate also showed an optimum of 0.3 and 0.5 mutations per daughter cell (Fig. 4H). While a higher mutation rate generates more high-affinity variants, it also increases the risk of mutating away before they have had time to divide, and produces more lethal mutations that reduce the number of progeny re-entering selection. Thus, optimal number of high-affinity plasma cells are generated when these processes are balanced at ∼0.3 mutations per daughter cell (Fig. S5G, H). The bias towards generating low-affinity variants also monotonically increases memory B-cell clonality (Fig. S5I).

We examined the combined effects of plasma cell differentiation and mutation rates on early (GC cycle 20) and late (GC cycle 40) high-affinity plasma cell counts (Fig. 4I). Optimal conditions for high affinity antibody production were stringent early on, peaking at 0.3 mutations per daughter cell and 3% plasma differentiation. However, after affinity maturation is established, more forgiving conditions allowed optimality across 0.1 to 0.5 mutations per daughter cell and 3% to 10% differentiation rates. In contrast, total titer varied monotonically, as when parameters were individually varied (Fig. S5J).

Overall, our findings provide intuition for the experimental outcomes, and further predict that mutation and differentiation rates play important but non-monotonic roles in shaping affinity maturation dynamics through their impacts on high-affinity outliers. The optimal rates predicted by the model showed striking congruence with physiological estimates of ∼0.5 mutations per cell ^39,40^ and 5% differentiation ^23,49^ highlighting the model reliability as well as the fine-tuning of the GC for an effective plasma cell response.

## DISCUSSION

Here we report that the regulated fragility and stability of the epigenetic state of B-cells has a role in determining the efficiency of generating high-affinity antibodies. Previous studies indicated that B-cell epigenetic states are rendered fragile during stochastic sampling of selection signals in the light zone ^15^, while being stably maintained during rapid proliferative bursts in the dark zone ^18^. Scrambling the epigenetic state in the light zone allows some highest-affinity B-cell clones to escape differentiation into plasma cells, fueling further rounds of affinity maturation. Conversely, the stable epigenetic state during proliferative bursts ensures maximum expansion of these high-affinity clones. Our insights resolve a key conflict in the literature of whether the seemingly stochastic B-cell fate decision making ^15–17^ may undermine the effectiveness of Darwinian genetic selection described by classical clonal selection theory ^13^.

Epigenetic variability in a population confers known evolutionary benefits ^8^ leveraged for bet-hedging by diverse organisms including viruses ^50–52^, bacteria ^53,54^, yeast ^55^, and mammalian tumor cells ^7,56^. Heritability of epigenetic states was also shown to advantage genetic alleles in fluctuating environments ^57,58^. In other cases, a lack of heritability can generate asymmetric cell fates during development ^59,60^. In contrast, antibody affinity maturation is a developmental process involving selection of genetic variants for high affinity, but where affinity-defined fitness is not monotonic with the selected trait, as the highest-affinity cells terminate proliferation to become immune effector plasma cells ^61^. These features define a unique role for epigenetic variability: to balance the need for effector formation while ensuring rapid affinity maturation, the germinal center reaction employs two alternating phases of generating epigenetic variability (fragility) and maintaining the epigenetic state (stability). Further, the generation and maintenance of genetic and epigenetic variation is notably in anti-phase, as AID-mediated mutation occurs during the proliferative burst in the dark zone while the epigenetic state is stable, but not in the light zone during selection when the epigenetic state is affected by stochastic encounters with numerous cells and stimuli. This phase-structured coordination of epigenetic and genetic variation across selection and expansion enables the germinal center reaction to simultaneously optimize multiple, potentially competing, properties of an effective antibody response.

Our work also underscores the importance of genetic and epigenetic outliers rather than the average B-cell in driving affinity maturation. Outlier cells and rare events dominate diverse immune processes from antiviral gene expression during infection ^62^ to selection within the GC itself ^63–65^, hence a predictive understanding must identify functionally relevant outliers and distinguish them from noise. This may challenge data-driven statistical inference methods that converge on a mean or dominant signal ^66,67^, but highlight a role for mechanistic models that can track relevant outliers. The present model formulation accounts for outliers on the single dimension of fitness score combining affinity and epigenetic cell state. Future refinements could extend the framework to account for distinct molecular modules controlling survival, proliferation, and differentiation propensities, whose outlier distributions can interact in complex ways to shape affinity maturation.

The mechanistic model reveals non-monotonic relationships between molecular regulatory parameters and affinity maturation, identifying optimal values that maximize antibody affinity gain or plasma cell production. Strikingly, these predicted optima align with physiological measurements in the literature (Fig. 3H ^23,39,40,49^), indicating that the process is finely tuned to maximize affinity gain, despite a wide parameter range where it remains robust but less efficient.

Further, trade-offs are apparent among multiple aspects of the response, as improvements in speed come at the cost of plasma cell generation, and likewise between clonality and mutational depth. This suggests a Pareto front balancing the various properties of a vaccine response, and that interventions for improvement should target the goldilocks zone.

Though we formulated the mathematical model to gain fundamental insight into how epigenetic heterogeneity affects evolution of genetic antibody variants, our work suggests that the model also has forward predictive power, generating antibody response outcomes under molecular regulatory perturbations or for individuals with different genetic or health status. The mechanisms encoded in this knowledge-based model constrain its outputs and hence require far less training data for accurate prediction than purely data-driven models, with the potential to extend these predictions beyond the training data. Importantly, its interpretability allows identification of regulatory targets for potential pharmacological perturbation and clinical translation. We suggest that clinical big data refinements of the mechanistic model may enable prediction of how small molecule modifiers of signaling pathways can be used as adjuvants or co-treatments to improve outcomes in poorly responding patient cohorts, to advance personalized vaccine administration.

### Limitations of the study

Our findings are based on a mathematical model that incorporates a wealth of prior research in its underlying assumptions, structure, and parameter selection. The model’s reliability is strengthened by sensitivity analyses, which revealed relationships aligned with expectation, measured rates of regulatory parameters, and perturbation studies using mutants to alter the regulatory network controlling B-cell fate decisions. We specifically addressed variability in B-cell fate decision-making in response to identical signals, by abstracting the biological complexities of other GC processes. While other models have depicted stochastic interactions with antigen-presenting follicular dendritic cells and T-follicular helper cells during B-cell selection ^28,68^, we reduced these to a dynamic affinity threshold consistent with current paradigms ^32^. We also assumed perfect affinity-dependent selection to isolate the effect of cell fate variability, justified by the sequential and largely independent nature of the two processes. We disregarded epitope-specificity of B-cell clones analogous to Brainbow-labeling studies of GC clonal dominance ^30^, as well as mutation biases to assume uniform exploration of genetic variants. These strategic abstractions enabled us to focus on the fundamental question of how B-cell epigenetic variability impacts affinity maturation.

## RESOURCE AVAILABILITY

### Lead contact

Further information and requests for resources and reagents should be directed to and will be fulfilled by the lead contact, Alexander Hoffmann.

### Data and code availability

- All modeling and analysis code are available on the Hoffmann Lab GitHub at the URL https://github.com/signalingsystemslab/GC_evolution_model.
- Raw data (FCS files, ELISA readouts) are also made available on the same GitHub repository.
- Any additional information required to reanalyze the data reported in this paper is available from the lead contact upon request.

## ACKNOWLEDGMENTS

We thank the members of the Hoffmann lab for valuable discussions and feedback on the manuscript, particularly Helen Huang, Chengyuan Li, Xiaolu Guo, Patrick Yuan, and Joseph Schirle as well as Roy Wollman and Eric Deeds for critical feedback and/or review of our manuscript. We acknowledge funding from NIH R01AI132731 and R01AI127867 to AH, the JSMF and Damon Runyon Quantitative Biology (DRQ11-21) Postdoctoral Fellowships to HVN, and a UCLA Undergraduate Research Scholarship to VK.

## AUTHOR CONTRIBUTIONS

Conceptualization, A.H. and H.V.N.; methodology, M.Y.X. and H.V.N.; investigation, M.Y.X., H.V.N., V.K., and T.W.; visualization, M.Y.X.; writing—original draft, M.Y.X., H.V.N., and A.H.; writing—review & editing, M.Y.X., H.V.N., V.K., T.W., and A.H.; funding acquisition, A.H.; software, M.Y.X.; supervision, A.H.

## DECLARATION OF GENERATIVE AI AND AI-ASSISTED TECHNOLOGIES

During the preparation of this work, the authors did not use any generative AI or AI-assisted technologies. The authors take full responsibility for the content of the publication.

## SUPPLEMENTAL FIGURES

**Figure S1.**
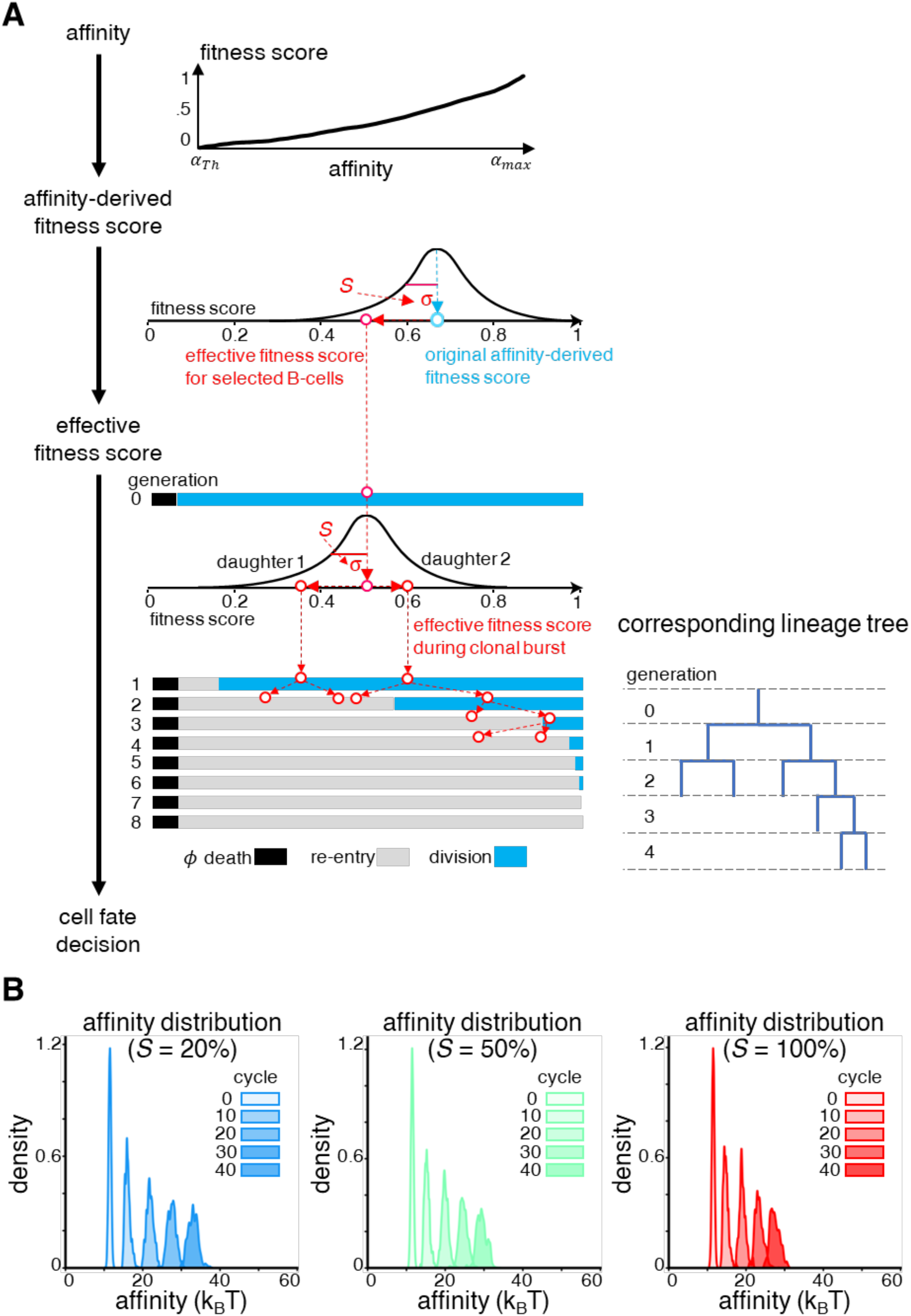
Stochastic B-cell fate decisions during affinity maturation. **(A)** Schematic showing the transformation of B-cell affinity into stochastic cell fate decisions that generate a proliferative lineage. ***(Top)*** Affinity is transformed into a fitness score representing the ranking of each B-cell within the selected population, as a proxy for how much T-dependent signaling is likely to have been received. ***(Middle)*** The affinity-derived fitness score is transformed into an effective fitness score for each cell, by drawing from a normal distribution centered at the original affinity-dependent fitness score, with a spread determined by the degree of stochasticity in cell fate decisions. ***(Bottom left)*** When the B-cell is assigned an effective fitness score, it picks the cell fate associated with that fitness score, in this case division of the selected B-cell. In the subsequent generations, each daughter B-cell is independently assigned a new effective fitness score, drawn from a normal distribution centered around the parent’s effective fitness score and with the same stochasticity-dependent spread. At each generation, the daughter cells undergo the fate decision determined by their newly determined effective fitness score. This independence between all progeny is consistent with prior probabilistic models of B-cell proliferative bursts, such as the Cyton model. ***(Bottom right)*** The cumulative set of fate decisions for all resulting progeny generate a lineage tree for the proliferative clonal burst arising from the selected B-cell during that cycle. **(B)** Kernel density plots showing the distribution of affinities of all B-cells re-entering the subsequent selection cycle, at regular intervals during affinity maturation for increasing values of stochasticity. While the starting affinity distributions at cycle 0 are narrow and identical, the distributions broaden over time. The tail of high-affinity outliers at cycle 40 is diminished as stochasticity rises, being present at 20% but mostly absent at 100% stochasticity.

**Figure S2.**
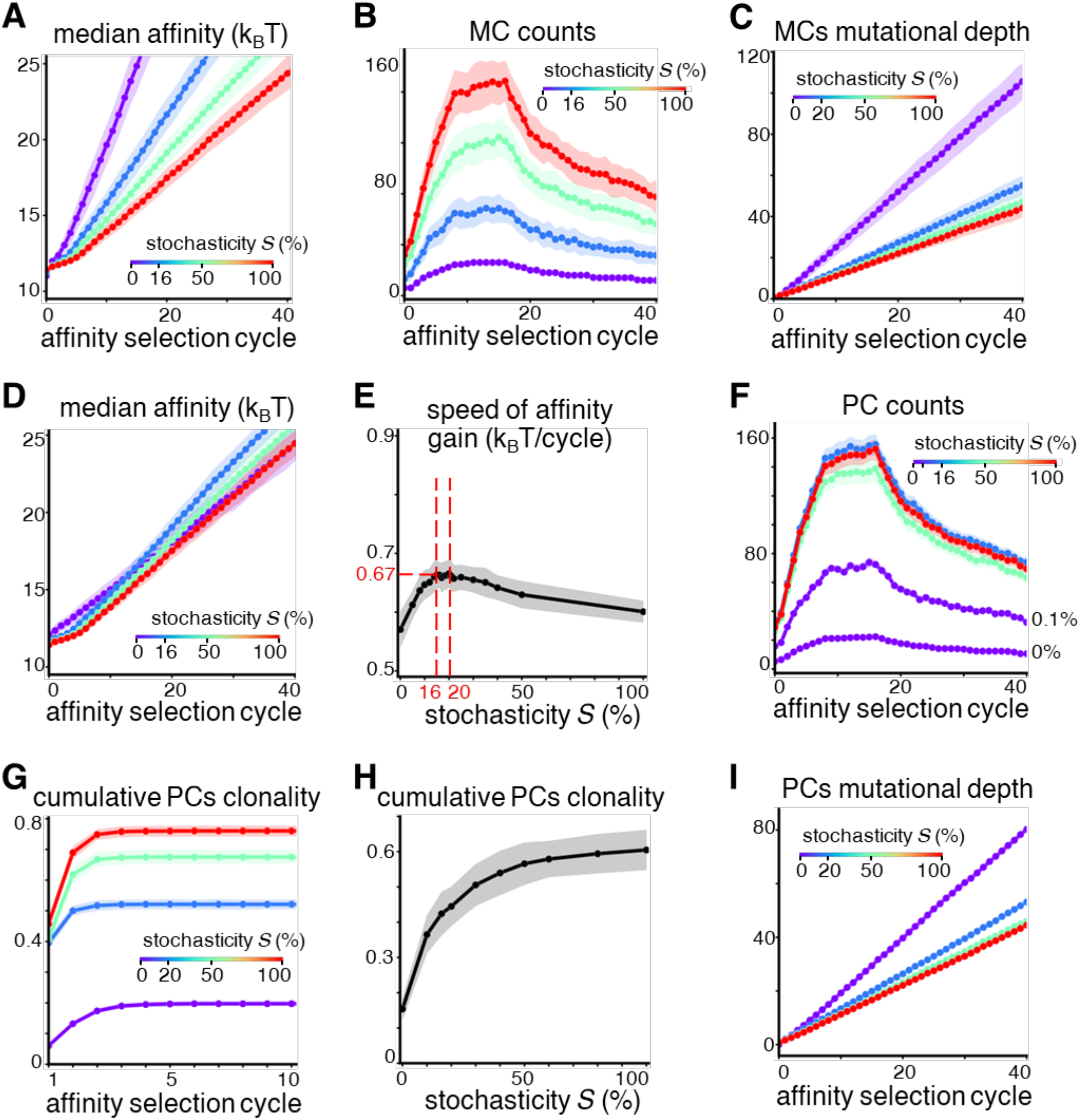
Multi-dimensional properties of plasma and memory B-cell effector compartments characterizing affinity maturation outcomes with varying stochasticity. **(A-C)** Line plots of **(A)** the median affinity, **(B)** the total number, and **(C)** the mutational depth (defined as number of mutations accumulated relative to the initial founder B-cell) of memory B-cells produced at each cycle of affinity maturation, for varying degrees of stochasticity in cell fate decisions. **(D-F)** Line plots of **(D)** the median affinity and **(E)** the speed of affinity gain (defined as the increase in median affinity relative to the previous selection cycle) of plasma cells, and **(F)** the total counts of plasma cells produced, at each cycle of affinity maturation for varying degrees of stochasticity in cell fate decisions. The vertical red lines in **(E)** indicate the region where speed of affinity gain is highest, between 16% to 20% stochasticity. **(G-I)** Line plots showing **(G, H)** cumulative clonal diversity of plasma cells and **(I)** mutational depth for plasma cells at each cycle of affinity maturation for varying degrees of stochasticity in cell fate decisions. In all cases, points indicate the median and shaded regions indicate the 95% confidence interval across 100 simulation runs.

**Figure S3.**
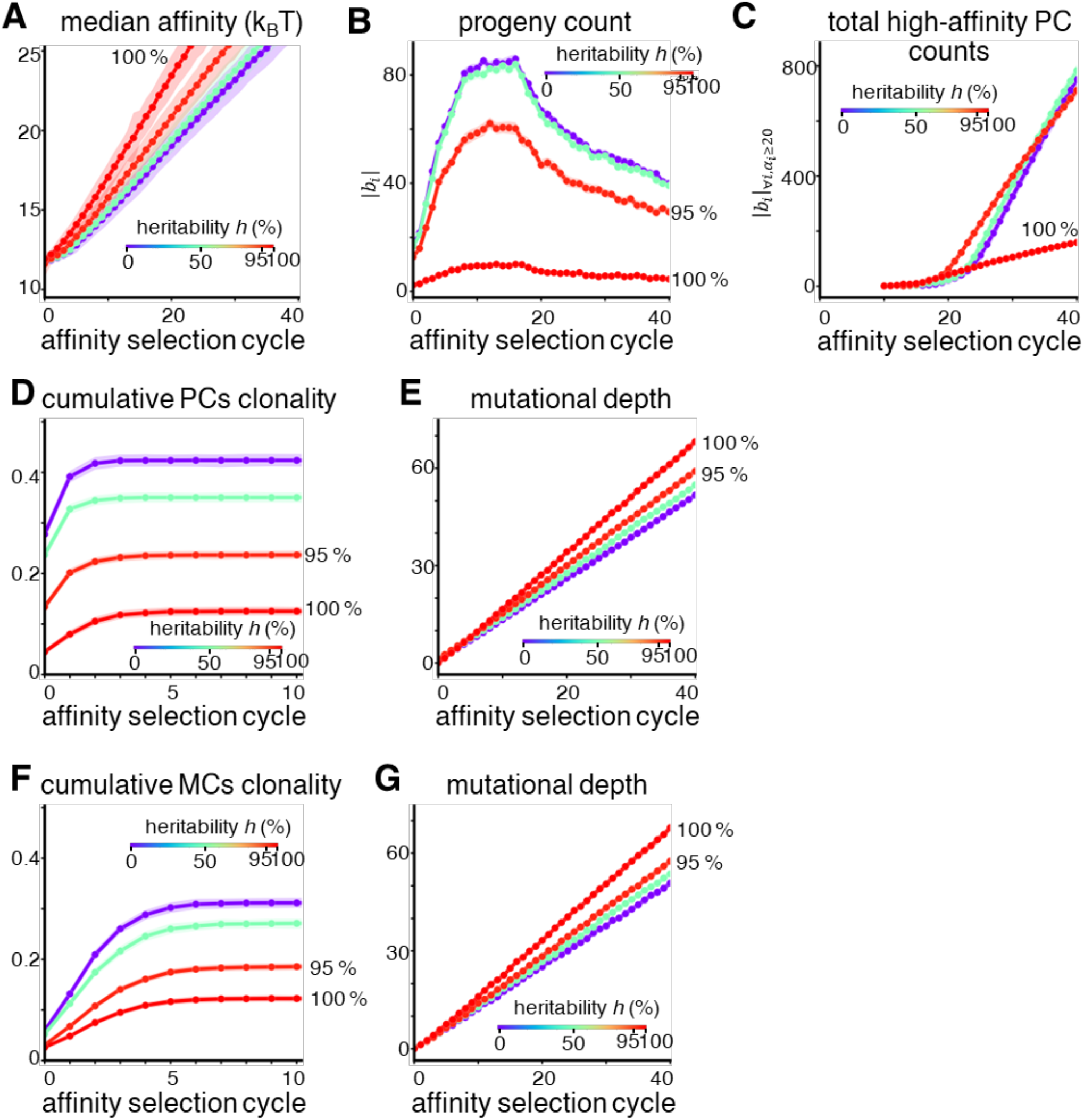
Multi-dimensional properties of plasma and memory B-cell effector compartments characterizing affinity maturation outcomes with varying heritability. **(A)** Line plots showing the variation of median plasma cell affinity, **(B)** median progeny per B-cell across selection cycles in the germinal center, and **(C)** the total counts of plasma cells above the high-affinity threshold (20.7 *k*_*B*_*T* or *K*_*D*_ 1 nM), computed at optimal stochasticity (16%) and different values of heritability. **(D-E)** Line plots showing **(D)** clonality (defined as fraction of extant clones relative to number of founder B-cell clones seeding the germinal center at cycle 0) and **(E)** mutational depth (defined as number of mutations accumulated relative to the initial founder B-cell) for plasma cells, at optimal (16%) stochasticity and varying degrees of heritability. **(F, G)** Similar plots are shown for the memory B-cell compartment, assumed similar to the set of B-cells eliminated by death, since both represent removal of low-affinity B-cells from the affinity maturation process. In all cases, points indicate the median and shaded regions indicate the 95% confidence interval across 100 simulation runs.

**Figure S4.**
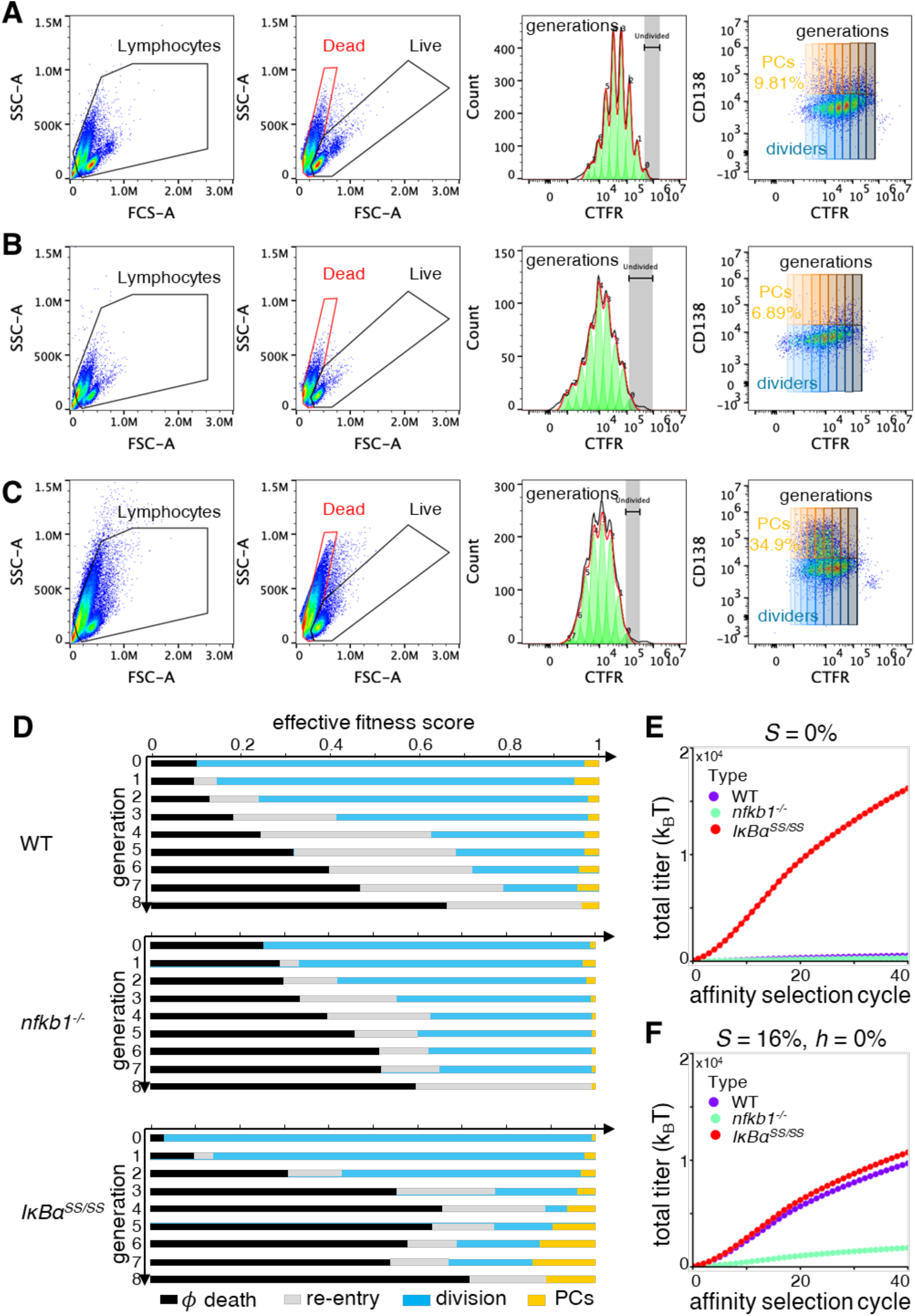
Measurement of probabilistic B-cell fate maps for different mouse mutant genotypes using in vitro flow cytometry data. **(A-C)** Raw flow cytometry plots of B-cells cultured for 120 hours, showing gating strategies to identify cells in different fates in **(A)** wild type, **(B)** *nfkb1*^-/-^, and **(C)** *IκBα*^SS/SS^ mice. Total lymphocytes are gated using forward scatter (FSC-A) and side scatter (SSC-A), with singlets identified using forward scatter area and height (plots not shown). Live and dead cell populations are separated based on differential clustering in the FSC-A vs SSC-A plot. Generations are identified using the intensity of Cell Trace Far Red (CTFR) stain, using the built-in proliferation module in FlowJo V10.0 applied to live cells. Plasma cells are identified by high expression levels of the marker CD138. The identified proliferation peaks are converted into gates, and applied uniformly to live, dead, and plasma cells, to separate all three populations by generation. For each genotype, at least 100,000 events representing 40% of total cells in culture were detected by flow cytometry, to increase statistical confidence in the measured proportions. **(D)** Probabilistic cell fate maps derived from the above *in vitro* flow cytometry data for wild type, *nfkb1*^-/-^, and *IκBα*^SS/SS^ genotypes, showing the proportions of B-cells undergoing each fate (death, cyclic re-entry, division, plasma cell differentiation) in each generation. **(E, F)** Model-predicted binding antibody titers (estimated as the cumulative plasma cell affinities) in each of the three genotypes, based on probabilistic cell fate maps generated from the *in vitro* flow cytometry measurements, but with **(E)** 0% stochasticity corresponding to deterministic clonal selection theory and **(F)** 16% stochasticity and 0% heritability corresponding to the Cyton model. Each point represents the median titer value across 100 simulation runs.

**Figure S5.**
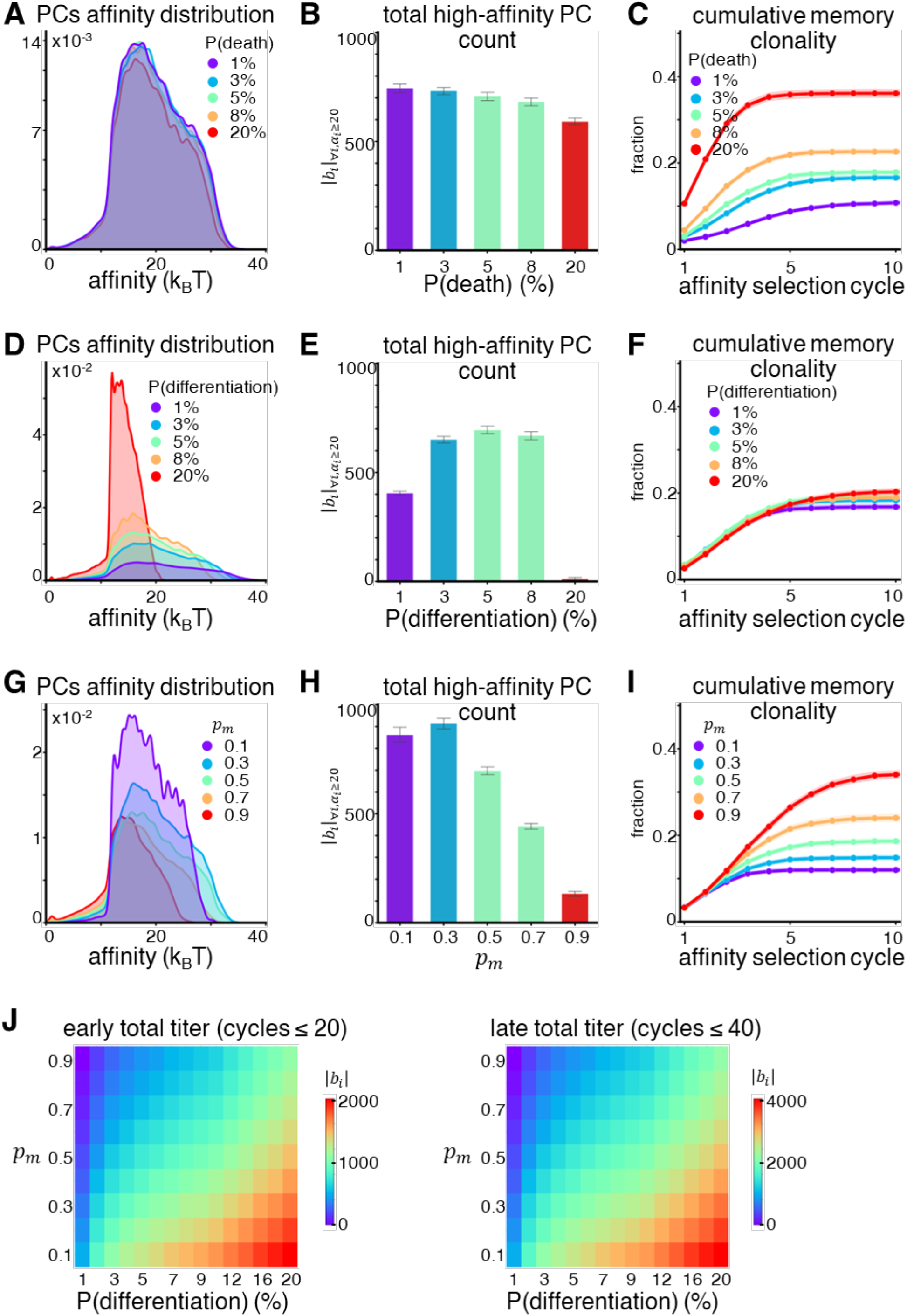
Properties of the two differentiated effector compartments, plasma cells and memory B-cells, as functions of key B-cell fate decision parameters during affinity maturation. **(A)** Kernel density plot showing the cumulative distribution of plasma cell affinities with **(B)** bar charts of the total number of plasma cells above the high-affinity threshold at the 40^th^ cycle (where error bars indicate the 95% confidence interval of counts across 100 simulation runs), and **(C)** line plots showing how clonality of the memory B-cell compartment evolves over the first ten selection cycles (where points indicate the median and shaded regions indicate the 95% confidence interval across 100 simulation runs) at varying rates of cell death. These plots are also indicative of outcomes for memory B-cell differentiation, which is equivalent to cell death in removing low affinity B-cells from the affinity maturation process. **(D-I)** The same three properties are also shown for varying rates of **(D-F)** plasma cell differentiation and **(G-I)** B-cell receptor mutation. **(J)** Heat maps of total antibody binding titer (estimated as the product of the number of plasma cells and their binding affinities) in early cycles (left) and at later cycles (right), as a function of varying plasma differentiation and mutation rates.

## STAR★METHODS MODEL CONSTRUCTION

We describe the construction of a model of B-cell evolution within the germinal center, based on experimental parameters derived from the literature. The overall model consists of selection, fitness and mutation functions operating upon a population of B-cells. Each of these modules is described in detail within the following sections.

### Defining the B cell population undergoing evolution

As the core component of the model, we define the population of B cells undergoing evolution, denoted as the time-varying set *B*. During any given evolutionary cycle *t*, we define

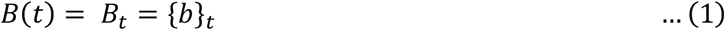

where *b* represents any B cell forming an element of the population set.

We characterize the population of B cells firstly by its size, i.e. number of elements in the set, as

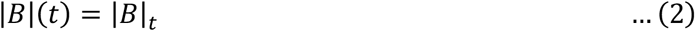

and secondly by its distribution of receptor affinities. The set of affinities *A* is defined as

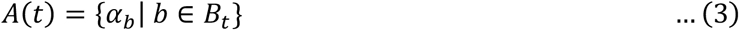

where *α*_*b*_ represents the affinity of any individual B cell *b* present in the population *B* at cycle *t*. We then allow this population of B cells to undergo 40 cycles of evolution, from *t* = 1 to *t* = 40, estimated to correspond to approximately 3 weeks of antigen response ^20^.

### Initialization of the model by setting B cell population parameters

At *t* = 0, we initialized the model with a population of 50 founder B-cells initiating the evolutionary cycle. This value was chosen from estimates of the number of clones seeding a typical germinal center, measured by Tas et al ^30^.

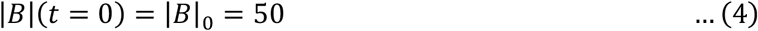

We assumed that the initial affinities of all the founders were normally distributed. We chose the mean of this normal distribution as a typical “low” affinity value, corresponding to an antigen-receptor dissociation constant *K*_*d*_ = 10 *μM* ^31^. This value can also be expressed on a linear scale, in terms of the free energy of binding Δ*G* between the antigen and receptor, using the relationship 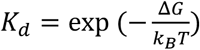 where *k*_*B*_ = 1.38 ∗ 10^−23^ *J*/*K* is the Boltzmann constant and *T* = 310*K* represents body temperature in a warm-blooded animal. Thus, in terms of relative energy units, the mean affinity of the founder cells 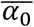 is fixed at 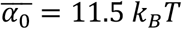 where 1 *k*_*B*_*T* = 4.3 × 10^−.0^ *J*.

Further, we assume that the spread in affinity between these founder B cells is small, such that 99% of all cells have affinities within the range of thermal fluctuations, i.e. within ± *k*_*B*_*T*. Thus, we choose 3σ_0_ = *k*_*B*_*T*, where σ_0_ is the standard deviation of the founders’ affinity distribution.

Overall, we assign the affinity of each founder B cell *α*_*b*_ by sampling values from a normal distribution with mean 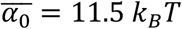 and standard deviation 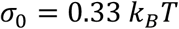, represented as

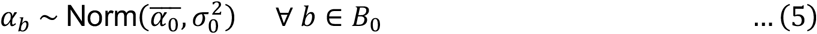

### Selection operating upon the B cell population

Selection of B-cells arises from their interactions with antigen and follicular helper T-cells, with high affinity as the phenotype being selected for. Without capturing the minutiae of interactions, we instead develop an abstraction of this process, allowing modulation of the stringency of selection. We assume that T cell help is the primary driver of selection ^69^, and is a limited resource partitioned across the B cell population based on their affinities. Thus, higher affinity B cells are more likely to receive T cell help, while lower affinity B cells receive none. We define the criterion for selection as a B cell having sufficient affinity to get T-cell help. In its simplest form, the probability of selection *p*_*Th*_ is given by:

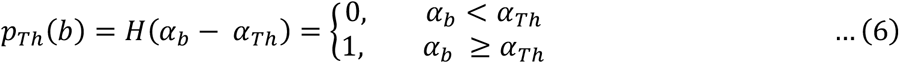

where *H* represents the Heaviside step function, such that only the B cells with affinities higher than a dynamic affinity threshold *α*_*Th*_ are selected while the rest are all eliminated.

We indirectly determine the value of *α*_*Th*_ at each cycle *t*, by prescribing the size of the population |*B*|_*t*_ using experimental observations of germinal center size over time ^34^. The value of *α*_*Th*_ is chosen such that the number of selected B cells above this affinity threshold matches this prescribed size. We then update the set *B* to get:

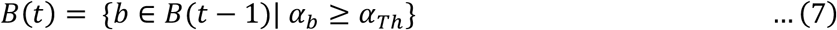

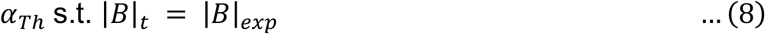

The selected cells in *B*(*t*) then undergo a cell fate decision between apoptosis, division, and plasma cell differentiation, with the progeny diversifying their receptor sequences through mutation, to complete the evolutionary cycle *t*.

### Cell fate decisions and fitness in the subset of selected B cells

For the set of B cells that receive T-cell help and are selected, we then map their affinities to cell fate decisions (i.e. phenotype to fitness), based on the classical clonal selection theory ^36^. The underlying assumption is that even among B cells that receive T cell help, the quantity of this help is partitioned based on their affinities. B cells on the lower end of the affinity distribution *A* receive only a little help and are more likely to die or divide a few times, while those on the higher end of *A* receive more help and are likely to expand further, or even differentiate into plasma cells at the highest affinities.

To model this, we first define a fitness score 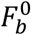 for each B cell, by transforming its absolute affinity *α*_*b*_ onto a common relative scale. We partition T cell help, and hence the fitness score, proportionally with affinity for each B cell, such that the fitness score corresponds to the rank of the B cell within the affinity distribution *A*. In mathematical terms, we map the affinity of each B cell *α*_*b*_ ∈ *A*[*α*_*min*_, *α*_*max*_] onto the fitness score 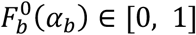 using the cumulative distribution function (CDF) of *A*, such that:

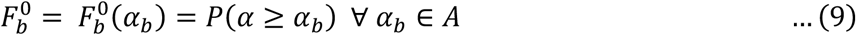

We then partition this common range of fitness scores into a map of B cell fate decisions, namely apoptosis, division, and plasma cell differentiation. We prescribe that fitness scores between [0, *p*_death_) correspond to apoptosis, since the experimentally measured apoptosis rate within germinal centers (excluding deaths due to both failed selection and lethal receptor mutation) is estimated around *p*_death_ = 5% ^37^. At the other end of the range, we prescribe that fitness scores between (*p*_differentiation_, 1] undergo plasma cell differentiation, as the plasma cell differentiation rate is also estimated to be around *p*_differentiation_ = 5% ^23,49^.

Fitness scores in the intervening range [0.05, 0.95] correspond to a clonal burst in the germinal center. Here, based on experimental observations ^38^, B cells undergo a burst of multiple divisions varying between 1 to 8, with an average of 3 divisions. We assume that following each division, the progeny independently decides whether to divide again and produce another generation, or to terminate the burst at that generation. Hence, we partition the interval of fitness scores according to terminal generation, by modeling the clonal burst as a binomial process over a range of generations, *G* ∈ [1, 8], having an expectation value 〈*G*〉 = 3. The probabilities of reaching each terminal generation cumulatively set their corresponding thresholds of fitness score. In other words, the probability *p*_*G*1_ of terminating the burst after a single division is mapped to fitness scores in the range [*p*_death_, *p*_death_ + *p*_*G*1_], a probability *p*_*G*2._ of dividing twice is mapped to (*p*_death_ + *p*_*G*1_, *p*_death_ + *p*_*G*1_ + *p*_*G*2._], and so on until the eighth and last generation which maps the interval (*p*_death_ + *p*_*G*1_ + ⋯ + *p*_*G*7_, *p*_differentiation_]. The resulting cell fate decision map is illustrated in Supplementary Figure 1A.

Finally, we assume that all B cell progeny that have reached their terminal generation re-enter the germinal center reaction for the subsequent cycle of evolution *t* + 1. To conveniently capture the decision between cyclic re-entry and division at each generation of a clonal burst, we treat the 1-D cell fate map of terminal generations across fitness scores as the linear projection of a 2-D cell fate map, where the second dimension now represents generation number. Thus, at each generation, the fitness score threshold corresponding to that terminal generation now marks the boundary between cyclic re-entry and division. Using the above example to illustrate, the first generation maps fitness scores in [*p*_death_, *p*_death_ + *p*_*G*1_] to re-entry and (*p*_death_ + *p*_*G*1_,*p*_differentiation_]to division, the second generation maps fitness scores in [*p*_death_, *p*_death_ + *p*_*G*1_ + *p*_*G*2._] to re-entry and (*p*_death_ + *p*_*G*1_ + *p*_*G*2_, *p*_differentiation_] to division, and so on until the eighth and last generation which maps the entire interval [*p*_death_, *p*_differentiation_] to cyclic re-entry. This 2-D expansion of the cell fate decision map across generations is also illustrated in Supplementary Figure 1B.

Following the assignment of cell fate decisions to each B cell *b*, we update the population *B*. Cells undergoing either apoptosis or plasma cell differentiation are removed from the population. The plasma cells are added to a separate set *P* (where we assume that there are no plasma cells prior to simulation, hence *P*(0) = ∅). The set of progeny {*b*_*G*_} from the terminal generation of a clonal burst are added to the population *B*_*t*_ and carried forward to the subsequent evolutionary cycle for selection. This modification of the population is represented as:

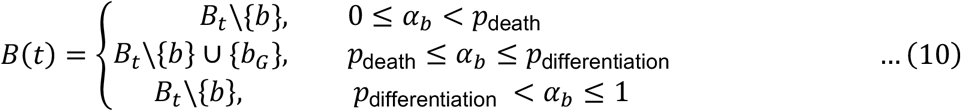

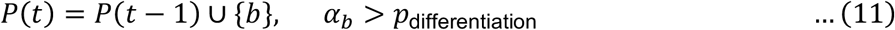

### Stochasticity and epigenetic heritability of cell fate decisions

Stochasticity in cell fate decision making may arise as a result of intrinsic noise in the intracellular molecular network regulating cell fates. To model this, we introduce a stochasticity parameter *S*, defined as the degree of decorrelation between the affinity-dependent initial fitness score 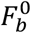 of a selected B cell, and the effective fitness scores *F*_*b*_ altered by noise at each generation of its proliferative lineage. We define *S* as:

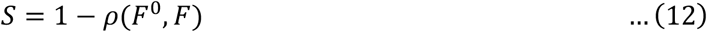

where 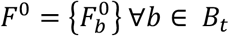 and *F* = {*F*_*b*_} ∀*b* ∈ *B*_*t*_ respectively represent the set of affinity-dependent and effective fitness scores for all B-cells in the population, and *ρ*(*F*^0^, *F*) is the Pearson correlation coefficient between them.

To do this for each 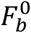, we draw the modified fitness score from a truncated normal distribution (restricted to the range [0,1] of possible fitness scores) around the mean value 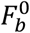. The spread of this distribution is given by a standard deviation σ_8_ which is determined by the desired degree of decorrelation *S*.

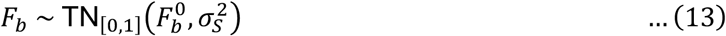

To approximate the value of σ_*s*_, we assume normality for the distributions of both *F*^0^ and *F*. The Pearson correlation coefficient between them is given by

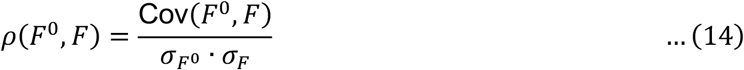

We assume 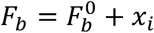 where *x*_*i*_ follows the normal distribution

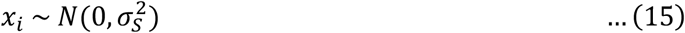

Thus, the mean of *F* remains unchanged since

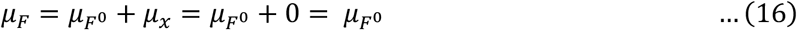

The variance of *F* is given by

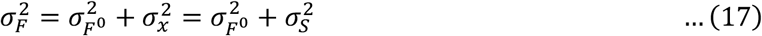

Hence the covariance between *F*^0^ and *F* can be simplified such that

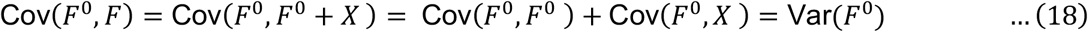

Since *F*^0^ and *X* are independent, Cov(*F*^0^, *X*) = 0.

Therefore, we can substitute this in Equation (14) for the correlation coefficient to get

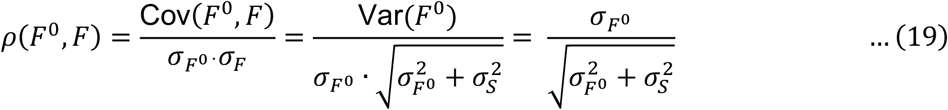

Therefore, we can determine σ_*s*_ based on the standard deviation 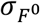 of B-cell fitness scores and the stochasticity parameter *S* = 1 − *ρ*(*F*^0^, *F*) by rearranging Equation (19) to get:

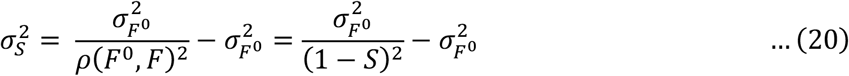

When cells enter a clonal burst, in the subsequent generations after the first, we also model a possible reduction in stochasticity due to epigenetic heritability among the progeny. To do this, we introduce a heritability parameter *h* to modify the stochasticity to get 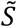:

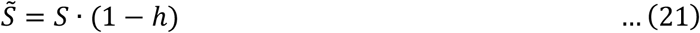

The value of *h* is chosen in the range *h* ∈ [0,1], where *h* = 1 represents perfect inheritance of cell fate propensities, and *h* = 0 indicates a complete lack of epigenetic correlation. After each cell division, we update the fitness scores for progeny *b*′ and *b*′′ based on their parent *b*, by modifying Equation (13) as:

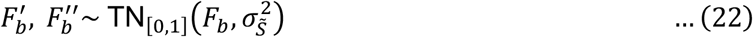

where *F*_*b*_′ and *F*_*b*_′′ represent the fitness score of the progeny after this redistribution. And calculate 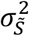 by modifying Equation (20) as:

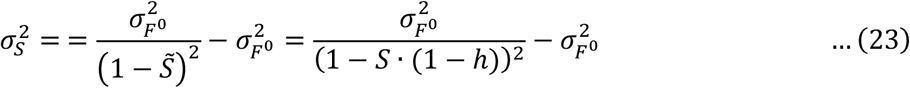

### Mutation in the subset of dividing B cells

B cells undergo somatic hypermutation (SHM), where the immunoglobulin locus undergoes point mutations that may alter the receptor sequence (genotype), and hence the receptor affinity (corresponding phenotype). This occurs only during divisions, and hence is confined to the subset of dividing B cells in each generation during a clonal burst. For each daughter cell, we assume that whether a mutation occurs or not can be modeled as a single Bernoulli trial with a mutation probability *p*_*m*_. We use a value of *p*_*m*_ = 0.5, derived based on estimates of the rate of SHM (∼1 mutation/1000 bp in each division ^39^) and the length of the immunoglobulin locus (∼600 bp ^40^).

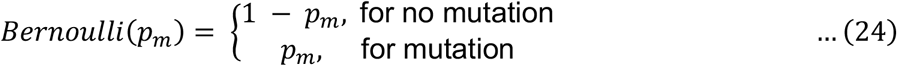

Once we determine that a daughter cell will undergo mutation, we next determine the impact of that mutation – whether it is synonymous, lethal, or changes the affinity of the cell. We assign the probabilities of each of these three outcomes to be 0.53, 0.28, and 0.19 respectively, based on theoretical estimates by Shannon and Mehr ^41^.

Daughter cells that undergo synonymous mutations are left unchanged in the population. In the case of lethal mutations, we remove those daughter cells from the population of B cells.

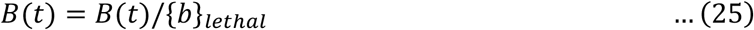

Finally, if we determine that a daughter cell will change its affinity due to mutation, we sample the change Δ*α* from a probability distribution determined previously ^42^, where only 5% of mutations are advantageous and 95% of them are deleterious. We update the affinity for any daughter cell *b*′ based on its parent *b* as:

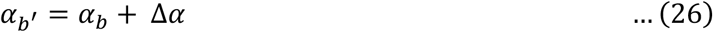

### Simulation details

The model code was written and run using Python version 3.11.4 on an Ubuntu server across 64 processing cores. To ensure the reproducibility of insights obtained from these probabilistic dynamics, we performed Monte Carlo simulations, averaging results across multiple simulation runs for each parameter set. We determined the minimum number required to ensure reproducibility as 50 runs, using a bootstrap method where we initially performed a large number of runs, down-sampled them in successively lower proportions, and set a criterion of median within 1% to assess similarity between original and down-sampled affinity distributions. In practice, we performed twice the number, i.e. 100 runs per parameter set, to provide maximum confidence that probabilistic trends were accurately and robustly captured without ambiguity.

### Table of all symbols and parameters used in the model equations

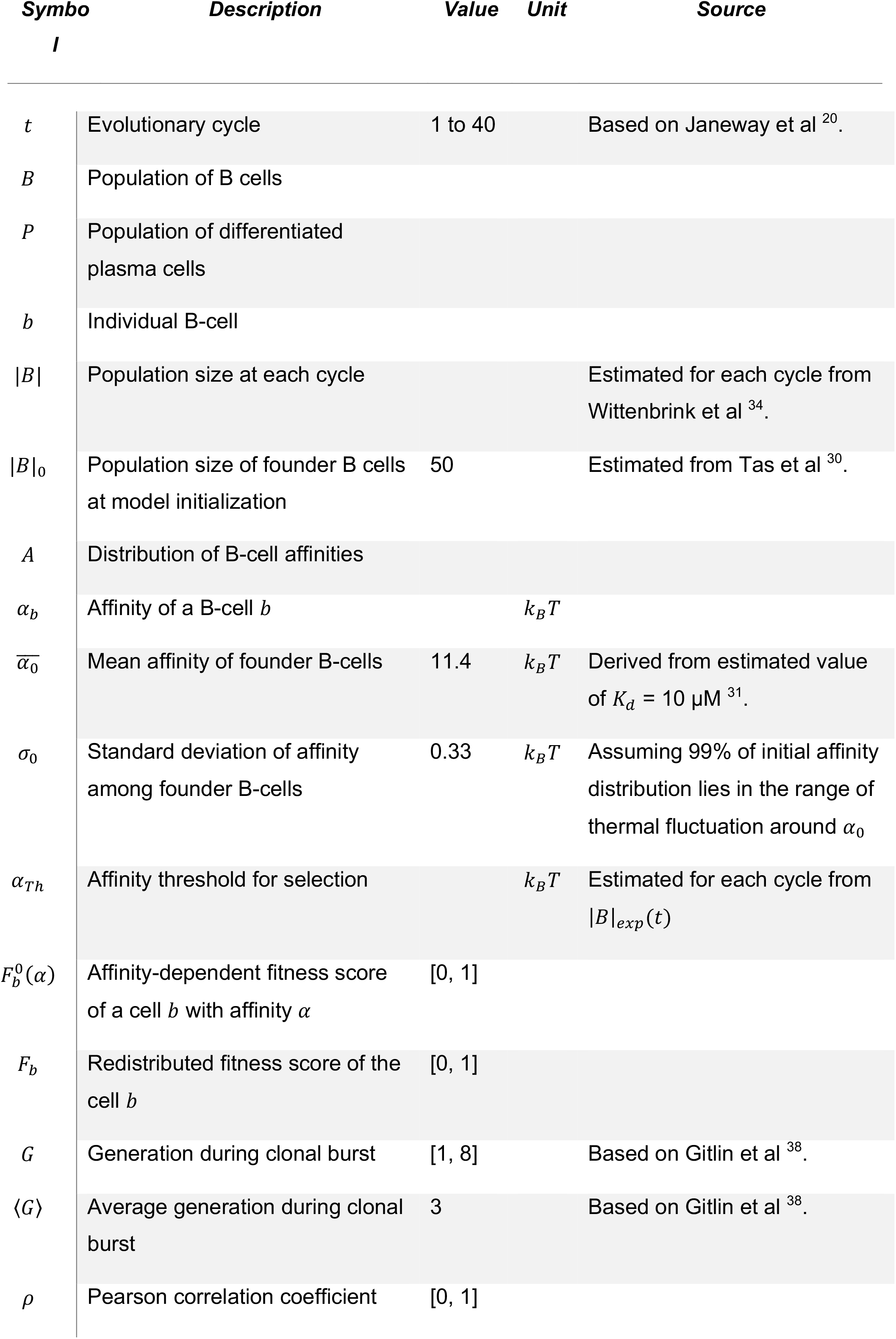
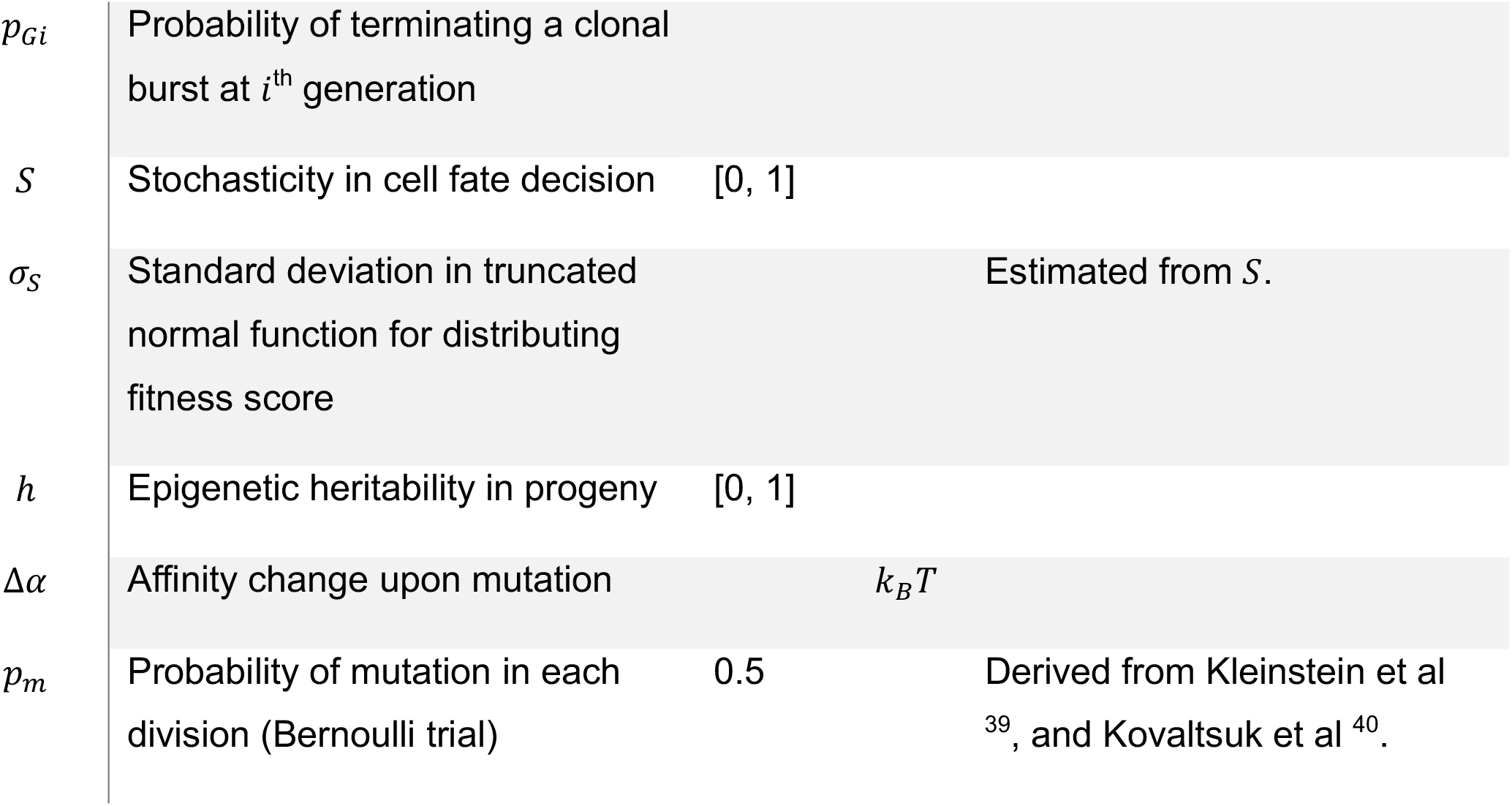

### KEY RESOURCES TABLE

| REAGENT OR RESOURCE | SOURCE | IDENTIFIER |
| --- | --- | --- |
| **Antibodies** |  |  |
| Rat monoclonal anti-human/mouse B220-APC | Biolegend | 103212; RRID: AB_312997 |
| Rat monoclonal anti-mouse CD138-PE (Clone 281-2) | Biolegend | 142504; RRID: AB_10916119 |
| Recombinant mouse monoclonal anti-NP [B1-8] standard 1mg/mL | Abcam | ab206535 |
| Goat anti-mouse polyclonal IgG-biotin | Abcam | ab6788; RRID: AB_954885 |
| **Chemicals, peptides, and recombinant proteins** |  |  |
| RPMI | Gibco | 11875-093 |
| CpG (ODN1668) | Invivogen | tlrl-1668-1 |
| CD154 (CD40 ligand) | Biolegend | 797406 |
| Murine recombinant BAFF/BlyS/TNFSF13B | R&D Systems | 2106-BF-010 |
| Murine recombinant IL-4 | PeproTech | 214-14 |
| Murine recombinant IL-21 | PeproTech | 210-21 |
| NP<sub>21</sub>-OVA | Biosearch Technologies | N-5051-10 |
| NP<sub>21-29</sub>-CGG | Santa Cruz Biotechnologies | sc-396209 |
| NP<sub>1-9</sub>-CGG | Santa Cruz Biotechnologies | sc-396187 |
| Alhydrogel | Invivogen | vac-alu-250 |
| **Critical commercial assays** |  |  |
| CellTrace™ Far Red Proliferation Kit | ThermoFisher Scientific | C34564 |
| Streptavidin-HRP | R&D Systems | DY998 |
| TMB color reagent for ELISA | R&D Systems | DY999B |
| **Experimental models: Cell lines** |  |  |
| NIH 3T3 cells | ATCC | CRL-1658 |
| **Experimental models: Organisms/strains** |  |  |
| Mouse: C57BL/6 | The Jackson Laboratory | JAX: 000664;<br>RRID:<br>IMSR_JAX:000664 |
| Mouse: C57BL/6: *nfkB1*<sup>-/-</sup> | The Jackson Laboratory | JAX: 006097;<br>RRID:<br>IMSR_JAX:006097 |
| Mouse: C57BL/6: *IkBα*<sup>SS/SS</sup> | The Jackson Laboratory | RRID:<br>IMSR_JAX:037800 |
| **Software and algorithms** |  |  |
| FlowJO V10.4.2 | FlowJO LLC | N/A |
| Python v3.7.164-bit base: conda | Anaconda v3.0 | N/A |

### EXPERIMENTAL MODEL AND SUBJECT DETAILS

#### Mice

Mice were maintained in environmental control facilities at the University of California, Los Angeles. Biological replicates were typically neither age-nor sex-matched, were not littermates, and often not co-housed, in order to determine whether differences between genotypes were indeed greater than differences between individuals within the same genotype. All mice used in experiments were 8-20 weeks old. The wild-type, *nfkb1*^*−/−* 70^ and IκBα^SS/SS 10^ mice were all on the C57BL/6 genetic background. Animal work was performed according to University of California, Los Angeles regulations under approved protocols.

## METHOD DETAILS

### Media and buffer compositions

*B cell media:* RPMI 1650 (Gibco) supplemented with 100 IU Penicillin, 100 µg/ml Streptomycin, 5 mM L-glutamine, 20 mM HEPES buffer, 1mM MEM non-essential amino acids, 1 mM sodium pyruvate, 10% fetal bovine serum (FBS), and 55 μM 2-Mercaptoethanol.

*MACS buffer:* Phosphate buffered saline (pH 7.4), 2% FBS, 100 IU Penicillin, 100 µg/ml Streptomycin.

*ELISA coating buffer:* Phosphate buffered saline (pH 7.4).

*ELISA sample dilution buffer:* Phosphate buffered saline (pH 7.4), 0.1% bovine serum albumin (BSA), 0.025% Tween-20.

*ELISA blocking buffer:* Phosphate buffered saline (pH 7.4), 1% BSA.

*ELISA wash buffer:* Phosphate buffered saline (pH 7.4), 0.1% BSA, 0.5% Tween-20.

### B cell isolation

Spleens were harvested from 8 to 20-week-old male and female C57BL/6 wild type, *nfkb1*^−/−^, and IκBα^*SS/SS*^ mice. For negative selection of B-cells, homogenized splenocytes were incubated with anti-CD43 magnetic beads for 15 min at 4-8 °C, washed with MACS buffer and passed through an LS column (Miltenyi Biotech). The purity of B cells was assessed at >98% based on B220 staining as described previously ^18^. Follicular B-cells were then further purified from the enriched B-cell population by positive selection, by labeling with anti-CD23 microbeads (Miltenyi Biotech) for 15 min at 4-8°C, followed by washing with MACS buffer and elution from an LS column. The purity of follicular B-cells was assessed at >99% using B220^+^ CD21^mid^ CD23^hi^ staining, distinguished from marginal zone (B220^+^ CD21^hi^ CD23^lo^) B-cells by flow cytometry.

### B cell in vitro culture

We developed a B-cell co-culture system to mimic germinal center follicles as closely as possible, based on prior work ^71^, while still retaining control over the quantity of various stimuli available to each B-cell. Briefly, NIH 3T3 fibroblasts were used to mimic the presence of stromal cells providing mechanical cues to B-cells, plated at a starting density of 10,000 per well in the 48-well plate and allowed to adhere for ∼8 hours prior to seeding follicular B-cells. Pure follicular B-cells were seeded at a density of 100,000 starting cells per well in a 48-well plate and cultured for 6 days in fresh RPMI-based media at 37 ºC and 5% CO_2_. Cells were stimulated with 50 ng/mL BAFF in solution to maintain survival. T-dependent stimuli were mimicked by adding 100 ng/mL CD154 (CD40 ligand) as a soluble peptide fragment, along with the cytokines 2 ng/mL IL-4 to promote proliferation and 1 ng/mL IL-21 to promote plasma differentiation. We determined these by first evaluating a range of doses for each stimulus to identify their lower non-responding and upper saturating doses, choosing their geometric mean as the representative dose in culture. B-cells were harvested at 120 hours to quantify proportions of various cell fates by flow cytometry.

### Measurement of generation-specific B cell fate decisions by flow cytometry

Immediately following isolation and prior to culture, WT, *nfkb1*^−/−^ and IκBα^*SS/SS*^ B cells were stained with Cell Trace Far Red (CTFR) using the CellTrace Far Red Cell Proliferation Kit (ThermoFisher Scientific, # C34564) as described by the manufacturer protocol. Briefly, 2M cells were resuspended in 1 mL RT PBS and incubated with 1 μL CTFR for 25 min at RT with rotation. Cells were washed by centrifuging, resuspending in 1 mL RPMI with 10% FBS, and incubating until continuous mixing for 10 min at RT. The washing steps were repeated a total of 3 times. CTFR labeled cells were cultured for 120 hrs as described above. B-cells were harvested by pipetting gently to resuspend them in the culture media without detaching the adherent 3T3 cells, collecting them in the supernatant, washing in PBS, and staining with 1:1000 anti-CD138-PE diluted in PBS with 15 minutes incubation for plasma cell identification. Samples were acquired on the CytoFlex flow cytometer (CytoFlex, Beckman Coulter). The cells were gated based on forward scatter (FSC) and side scatter (SSC) to identify live single cells. Doublets were excluded from the analysis using FSC area and height. Cell generation numbers were defined based on dilution of CTFR, and plasma cells identified as CD138^hi^. Further quantitative analyses to estimate cell numbers per generation undergoing each fate were performed as described below.

### Immunization and serum collection in live mice

Mice were immunized with the model antigen NP_21_-OVA (Biosearch Technologies) adsorbed on the adjuvant Alhydrogel 2% (Invivogen). A 5 mg/mL stock of NP_21_-OVA and 10% Alhydrogel were mixed in a 1:1 volumetric ratio by pipetting vigorously for 10 minutes. 8-to 20-week-old male and female C57BL/6 wild type, *nfkb1*^−/−^, and IκBα^*SS/SS*^ mice were immunized subcutaneously in the left hind footpad with 30 µL of the mixture using 31G insulin syringes, corresponding to 30 µg NP_21_-OVA and 60µg adjuvant for each mouse as per established protocols ^72^. Only a single priming dose was administered to monitor affinity maturation, as the computational model does not explicitly account for recruitment of memory cells in a boosted response. About 50 µL blood was collected retro-orbitally from each mouse in non-heparinized glass capillaries at 0-, 5-, 8-, 11-, 15-, and 18-days post immunization, alternating between left and right eyes to minimize injury. Blood samples were placed at room temperature for at least 3 hours to initiate clotting, maintained overnight at 4 ºC, centrifuged at 500 rcf for 5 minutes to separate clot from serum with minimum hemolysis, after which about 20 µL supernatant serum was collected. Centrifugation and supernatant collection were repeated to fully remove all debris, and purified serum samples were stored at -20 ºC until antibody content was assessed by ELISA.

### Measurement of serum antibody by ELISA

ELISA protocols were adapted from established protocols (R&D Systems). Total antibody quantity was estimated by binding against NP_25_ oligomers and high-affinity fraction by binding against NP_5_ oligomers, with both oligomers conjugated to chicken gamma globulin (CGG) as a neutral carrier distinct from the conjugate immunogen ovalbumin (OVA). ELISA plates were prepared for capture by coating with 1 µg/mL capture reagent (NP_21-29_-CGG or NP_1-9_-CGG, Santa Cruz Biotechnologies) diluted in PBS. 50 µL coating reagent was applied to each well of a 96-well half-area plate and incubated overnight at 4 ºC. Coated plates were washed according to a standard wash protocol of 3 times in ELISA wash buffer with 3 minutes of shaking each time, once more without shaking, and finally once in distilled water to remove excess coating reagent. All subsequent incubations were done at room temperature with continuous shaking for 1 hour unless specified otherwise. Plates were blocked in 200 µL ELISA blocking buffer before washing. Samples were diluted to concentrations between 1:10,000 and 1:100,000 in ELISA sample dilution buffer, 50 µL each applied to sample wells and incubated. Recombinant monoclonal B1-8 antibody (1 mg/mL stock) with known high affinity against NP was used to prepare standards at five serial dilutions between 1 µg/mL and 1 ng/mL, with anti-tubulin as a negative control, and ELISA sample dilution buffer as blanks. Samples were washed off and incubated with 50 µL/well detection antibody (goat anti-mouse IgG-biotin) diluted to a working concentration of 0.5 µg/mL in ELISA sample diluent buffer. After washing again, plates were incubated for 20 minutes with 10 µg/mL streptavidin-HRP (R&D Systems) and washed to remove all unbound substrates. 50 µL/well of TMB color reagent (R&D Systems) was added to each well and incubated for 3-8 minutes until blue color developed. The reaction was stopped by adding 25 µL of 2N H_2_SO_4_ and absorbances measured on a plate reader (Biotek Gen 5.1.1) at 450 nm for signal and 570 nm for correction. Blank wells were averaged and corrected absorbances calculated for each sample using the formula (Measured450 – Blank450) – (Measured570 – Blank570). Raw absorbance values in the linear portion of the standard curve were converted to absolute concentrations in mg/mL, and fractions of high affinity antibody estimated as the ratio of antibody bound to NP_5_ over that bound to NP_25_.

## QUANTIFICATION AND STATISTICAL ANALYSIS

### Fitting stochasticity and heritability parameters to experimental measurements

To fit the stochasticity parameter, we utilized experimental measurements from Mitchell et al ^18^ by comparing the terminal generation distribution between experimental data and first-cycle simulation results using the Mann-Whitney U test. Additionally, we assessed lineage symmetry by applying a t-test to compare experimental lineage measurements with first-cycle simulation results. Both statistical scores were normalized to a range of -1 to 1, and a combined fitting score was computed as the average of both, to evaluate agreement between experiment and simulation. We conducted 100 Monte Carlo simulations using bootstrap resampling with 50 random samples per iteration. This process was repeated 300 times to obtain an average combined fitting score for each pair of stochasticity and heritability parameter values. Finally, we varied both stochasticity and heritability parameters to generate a heatmap illustrating the parameter space and its overall fitting.

### Flow cytometry analysis to generate cell fate maps

Flow cytometry data were analyzed using FlowJo V10.0, with gating applied to identify lymphocytes, live lymphocytes, dead lymphocytes, and plasma cells (Fig. S5A). The proliferation analysis tool was used to track cell generations and calculate population sizes for live, dead, and plasma cells. Target cell fate propensities were then derived using a set of mathematical equations.

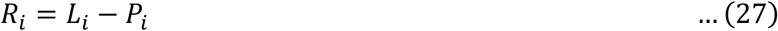

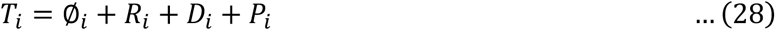

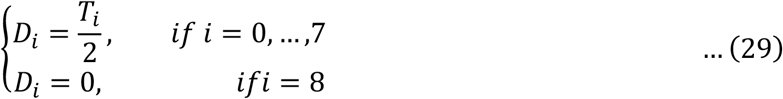

where *i* ∈ [0,8] represent B-cell generations during clonal burst, *R* represents re-entering cells, *L* represents live cells, *P* represents plasma cells, *T* represents total cells, ∅ represents dead cells, and *D* represents dividing cells. This is based on the logic that the total number of cells in a subsequent generation must be twice the number of dividing cells in the previous generation, with the boundary condition that all cells in the terminal generation undergo either death, plasma differentiation, or cyclic re-entry. We constructed the expanded 2D-fate map as follows:

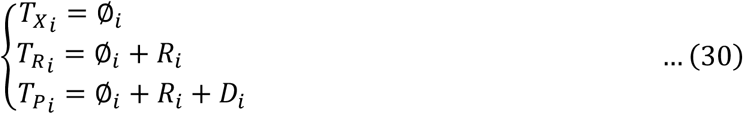

